# A functional map of CDK-drug interactions at single amino acid resolution

**DOI:** 10.1101/2025.11.02.685764

**Authors:** Samuel I. Gould, Jonuelle Acosta, Luca Boscolo Bielo, Alinur Jaboldinov, Grace A. Johnson, Manuel E. Contreras, Alexandra N. Wuest, Emmanuel Brémont, Mohammed A. Toure, Yichen Xiang, Zuzanna Kozicka, Michael T. Hemann, Angela N. Koehler, Sarat Chandarlapaty, Francisco J. Sánchez-Rivera

## Abstract

Proteins that drive or support human disease phenotypes are attractive molecular targets for precision therapy, yet most are nominated by knockout studies and then targeted with drugs that inhibit core catalytic pockets. These strategies cannot resolve which residues are essential, whether non-catalytic sites offer better selectivity or potency, or identify on-target resistance mechanisms. We introduce a framework that integrates precision genome editing, mechanistically diverse therapeutics, and computational sequence-structure-function analysis to map protein essentiality and potential druggability at single amino acid resolution. Applying this framework across 9 cyclin-dependent kinases (CDKs) and 15 cancer therapeutics—including ATP-competitive inhibitors, PROTACs, and molecular glue degraders—we identify shared and CDK-specific residues critical for cell fitness and drug response, including known resistance mutations and dozens of new variants. The resulting functional maps resolve residue- and mechanism-specific differences in the resistance spectra among agents targeting the same protein. We show that this iterative strategy can also uncover higher order interactions by performing intra- and extragenic epistasis screens to identify residues that mediate on-target and within-family cell fitness and drug resistance. Finally, we find evidence of novel *CDK6* mutations in breast cancer patients and concordance between experimental and clinical correlates of response to CDK4/6 inhibitors. By mapping residue-level essentiality and forecasting therapy resistance mutations, target-drug interaction maps could inform clinical treatment and guide design of more selective therapeutic molecules.

## Introduction

The clinical success of targeted cancer therapies like imatinib, trastuzumab, and gefitinib led to a major paradigm shift in oncology drug development to focus on treating cancer patients with molecular precision (*1*–*5*). This motivated the development of small molecules capable of inhibiting or modulating the activity of wild type (WT) or mutant cancer-associated proteins, which remains the subject of active research and drug development efforts worldwide. A large fraction of these therapies target kinases due to the central role that these proteins play in fundamental aspects of cancer cell signaling and proliferation (*6*). Oncogenic proteins with different enzymatic activities or non-catalytic molecular functions have also been identified and successfully drugged in the clinic (*7*), highlighting the need to understand the mechanistic diversity of cancer dependencies at a molecular level.

Advances in large-scale RNA interference and CRISPR-based functional genomics over the last two decades have streamlined the identification of cancer drivers, dependencies, and determinants of therapeutic response across many types of human malignancies (*8*–*10*). This pan-cancer dependency mapping can provide a rational framework to nominate therapeutic targets that may be effective across multiple cancer types or in defined genetic subgroups. In parallel, chemical biology approaches have expanded the universe of druggable targets and redefined cancer therapeutic development. Targeted protein degradation strategies have emerged as a powerful modality to target molecularly-diverse proteins, including factors long considered to be ‘undruggable’ by conventional inhibitors. This includes proteolysis-targeting chimeras (PROTACs) that recruit an E3 ubiquitin ligase to a target protein, and molecular glue degraders that induce neomorphic interactions between E3 ligases and target proteins (*11*).

Within this landscape, cyclin-dependent kinases (CDKs) have reemerged as critical therapeutic targets in oncology. CDKs are master regulators of cell cycle progression and transcription that are frequently co-opted by cancer cells to drive tumor growth and sustain oncogenic transcriptional programs (*12*, *13*). Notably, inhibitors of CDK4 and CDK6 recently achieved breakthrough therapeutic success in patients suffering from hormone receptor-positive breast cancer, leading to a significant extension of progression-free survival when combined with standard-of-care endocrine therapies (*14*–*22*). Clinical validation of CDK-targeting drugs has sparked interest in exploiting the reliance of cancer cells on sustained CDK activity, leading to the development of new inhibitors and dozens of clinical trials across solid tumors and hematological malignancies.

Despite the clinical success of targeted therapies against CDK4/6 and other cancer dependencies, treatment resistance is a universal obstacle to achieving durable remission and patient survival. Most targeted therapies are initially effective in reducing tumor burden, but treatment resistance inevitably emerges and renders these therapies ineffective. For instance, breast cancer patients almost invariably develop resistance to CDK inhibitors through diverse molecular mechanisms, including loss of *RB1*, upregulation of Cyclin E/CDK2 by *CCNE1* amplification or *c-MYC* overexpression, amplification of *AURKA*, or loss of *FAT1* (*23*–*27*). More broadly, many different mechanisms of treatment resistance have been observed, including but not limited to activation of compensatory signaling pathways, copy number amplification of the therapeutic target, upregulation of drug efflux pathways, cell state changes, and changes to the tumor microenvironment (*28*).

One common mechanism of resistance is mutation of the protein that is the target of the therapy of interest. These mutations often occur in the drug binding site, rendering the therapy ineffective through steric hindrance or reductions in binding affinity. There are many examples of this phenomenon across multiple therapies and targets, including in *EGFR, ALK, ABL1*, and *KRAS (29–32)*. Importantly, the identification of these resistance-conferring variants (hereafter referred to as resistance variants) has enabled the development of molecules capable of binding mutant forms of the target protein. This includes ponatinib and osimertinib, which are able to bind certain mutant *BCR-ABL* and *EGFR* proteins, respectively (*33*, *34*). However, despite the efficacy of these next-generation therapies, resistance can still emerge in the form of compound mutations that reduce binding affinity beyond therapeutically feasible levels of drug administration (*35*, *36*). The efficacy of any targeted therapy may therefore hinge on specific residues and surfaces within the target protein that influence drug binding, protein-protein interactions, and signal transduction regulation.

Given the critical role of drug resistance in determining the durability and efficacy of targeted therapies, the identification of resistance variants is of fundamental importance to oncology. Classically, resistance variants have been identified in the clinic after drugs are administered to patients; ideally, these variants would be identified long before drugs make it to the clinic. It is also possible that drug-resistant variants are already present at sub-clonal, undetectable levels within a primary tumor and only become apparent during the course of treatment or after therapy due to positive selection. These variants may not drive tumor progression *per se* or be of unknown functional significance and thus may be missed or ignored by clinical sequencing efforts. Though some studies have identified resistance variants and mechanisms against these agents, including CDK inhibitors (*26*, *37*–*42*), the full spectrum of resistance-conferring alterations remains to be defined. Prospective knowledge about these types of variants could guide clinical decision-making and rational design of improved therapeutic strategies.

We reasoned that high efficiency base editing sensor libraries (*43*–*47*) could be used to perform systematic tiling mutagenesis of drug targets (*43*, *48*–*59*) to identify essential protein residues and resolve residue- and mechanism-specific differences in the resistance spectra among agents targeting the same protein (**Figure 1a**). To test this concept, we generated base editing sensor tiling libraries covering 9 transcriptional or cell cycle CDKs and screened them across 15 cancer therapeutics, including ATP-competitive inhibitors, PROTACs, and molecular glue degraders, allowing us to pinpoint essential residues and mechanism-specific responses among these agents. By iterating on this strategy to perform intra- and extragenic epistasis screens, we identified CDK residues that mediate on-target and within-family cell fitness and drug resistance. Finally, we provide evidence of novel *CDK6* mutations in breast cancer patients and concordance between experimental and clinical correlates of response to CDK4/6 inhibitors. By mapping residue-level essentiality and forecasting therapy resistance mutations, this type of target-drug interaction mapping could inform clinical treatment and guide design of more selective therapeutic molecules.

**Figure 1.**
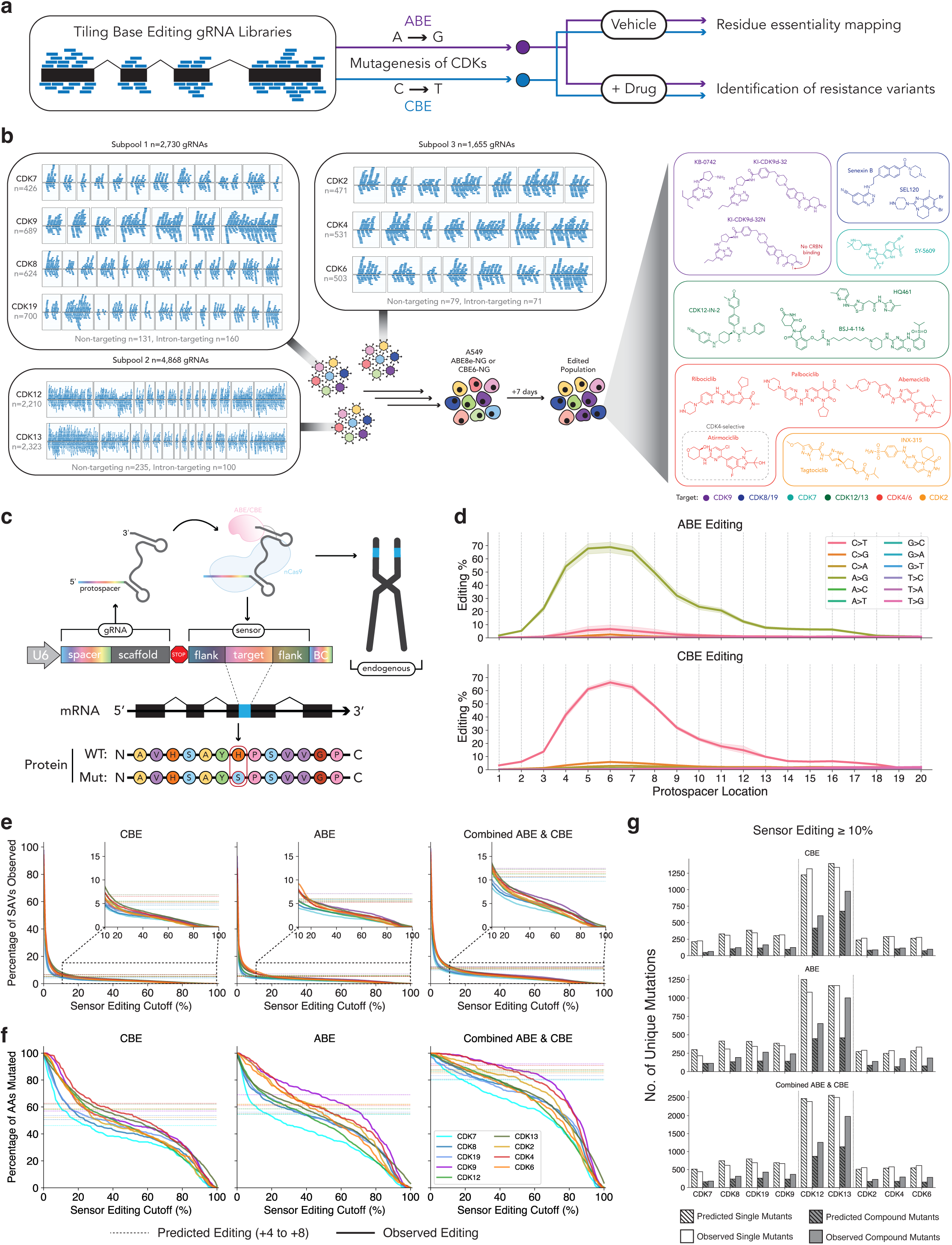
Base editing sensor tiling libraries enable multiplexed mutagenesis of the CDK family. **a)** Base editing libraries are designed to tile the CDS of target genes to enable mutagenesis with ABE and CBE. Once mutant pools are generated, cells are treated with targeted therapies to identify resistance variants, or treated with vehicle (DMSO) to map residue essentiality. **b)** Schematic of overall screening strategy. Each subpool was delivered in lentiviral form to either ABE8e- or CBE6-NG expressing A549 cells. After allowing editing to occur for 7 days, cells were treated with 1 of the 15 targeted therapies depicted in the figure for a period of ≥21 days. **c)** Diagram of the base editing sensor strategy. The sensor site encodes a copy of the endogenous target site and can be used to infer amino acid-level mutations for each gRNA. **d)** Quantification of ABE (top) and CBE (bottom) editing efficiency as a function of protospacer location. The shaded region represents a 95% confidence interval of the mean value across the quantified samples. **e)** Quantification of the percentage of SAVs observed (relative to the total possible number of SAVs) at sensor editing frequency cutoffs ranging from 0 to 100% for each gene. CBE (left), ABE (middle), and combined ABE and CBE (right). Dashed lines represent predicted editing based on complete mutagenesis of the +4 to +8 region of the protospacer of each gRNA. **f)** Same as (e), but quantifying the percentage of amino acids mutated as a function of sensor editing cutoff. **g)** Barplot quantifying the number of unique single and compound mutants for each gene observed in at least 10% of sensor reads for each gRNA, relative to the predicted number of mutations based on complete mutagenesis of the +4 to +8 region of the protospacer.

## Results

### Engineering & screening CDK variants for resistance against diverse targeted therapies

To map essential protein residues and identify resistance variants against distinct clinical-grade or investigational small molecules targeting CDKs, we designed base editing sensor tiling libraries for the human *CDK7, CDK8, CDK9, CDK12, CDK13,* and *CDK19* transcriptional CDKs, as well as *CDK2, CDK4,* and *CDK6* **(Figure 1a-c**; see **Methods** section). These libraries were rationally split into 3 distinct subpools to maximize screen representation and experimental throughput across many conditions and drugs. The subpools were separated based on CDK function: subpool 1 contains *CDK7, CDK8, CDK9,* and *CDK19*, which are involved in transcriptional initiation and elongation; subpool 2 contains *CDK12* and *CDK13*, the CDKs involved in the later stages of transcriptional elongation and termination; and subpool 3 contains *CDK2*, *CDK4*, and *CDK6*, corresponding to a subset of the cell cycle CDKs (**Figure 1b**).

We screened these libraries in A549 human lung adenocarcinoma cells expressing either ABE8e-NG (*60*) or CBE6-NG (*61*). Library-transduced cells were cultured for 7 days to ensure maximum editing followed by splitting into culture media with vehicle (DMSO) or 1 of 15 CDK-binding compounds that produce cytotoxic or cytostatic effects in A549 cells (**Figure 1a-b**, **Supplementary Figure 1, 2**). Subpool 1 (*CDK7/8/9/19*) variants were screened in the presence of (i) KB-0742 (*62*, *63*), an ATP-competitive CDK9 inhibitor, (ii) KI-CDK9d-32 (*64*), a CDK9 PROTAC, (iii) KI-CDK9d-32N (*64*), a compound with the same structure as KI-CDK9d-32, but lacking degradation activity, (iv) SY-5609 (*65*, *66*), a clinically tested ATP-competitive CDK7 inhibitor, and (v) SEL120 (*67*, *68*) and (vi) Senexin B (also known as BCD-115) (*69*, *70*), both of which are clinically tested CDK8/19 inhibitors. Subpool 2 (*CDK12/13*) variants were screened in the presence of (i) CDK12-IN-2 (*71*), an ATP-competitive CDK12/13 inhibitor, (ii) BSJ-4-116 (*39*), a CDK12/13 PROTAC, and (iii) HQ461 (*38*), a molecular glue degrader that mediates an interaction between CDK12/13 and DDB1 to facilitate ubiquitination of cyclin K. Subpool 3 (*CDK2/4/6*) variants were screened in the presence of three clinically approved CDK4/6 inhibitors, (i) Ribociclib (*72*), (ii) Palbociclib (*73*), and (iii) Abemaciclib (*17*), as well as (iv) Atirmociclib (*74*), a CDK4-selective, clinical-grade compound, and (v) INX-315 (*75*), and (vi) Tagtociclib (PF-07104091) (*76*, *77*), both of which are clinically tested CDK2 inhibitors. After 21 days of passaging and selection, genomic DNA was extracted from initial and final time-points to assess gRNA enrichment and depletion, as well as sensor editing to quantify base editing mutagenesis (see **Methods** section).

### Base editing sensor tiling screens enable mutagenesis of nearly all amino acids in the CDKs

Our screens use the base editing sensor architecture, where a copy of the endogenous target site is included adjacent to the gRNA sequence (*43*, *45*, *46*) **(Figure 1c)**. This architecture allows us to quantify the editing activity of each gRNA individually at the sensor site, which we and others have shown closely mimics the editing at cognate endogenous targets (*43*–*46*, *78*). This approach obviates the need to amplify and sequence each targeted exon, and provides a direct measurement of gRNA efficiency that would not be possible even if each exon was directly sequenced. Importantly, because the desired edits are not defined *a priori* in a base editing tiling screen, quantification of sensor editing enables the subsequent prediction of the effects of each edit at the amino acid level, where the resistance phenotypes are most likely to manifest **(Figure 1c).**

The window of maximal base editing activity is typically defined as the 5 nucleotide window spanning the +4 to +8 region of the protospacer (*43*, *60*, *61*, *79*). Quantifying the average sensor editing rate across thousands of gRNA-sensor pairs in >50 screen samples per editor, we confirmed that this +4 to +8 region is the region of maximal base editing activity (**Figure 1d**). However, we also observed considerable editing spanning the +2 to +17 region of the protospacer with both editors **(Figure 1d).** This result highlights the need to make empirical measurements of gRNA activity via the sensor system, as opposed to simply relying on the +4 to +8 maximal editing window heuristic, since edits often fall outside of this typical window.

Next, we mapped the base edits to the corresponding coding sequences of each targeted CDK to quantify the frequency of each amino acid substitution installed by each gRNA (**Figure 1e**; see **Methods**). This analysis allowed us to empirically determine the fraction of single amino acid variants (SAVs) that we could engineer out of the set of possible SAVs (20 possible substitutions at each amino acid position). For instance, at a frequency cutoff of 10%, we could engineer ∼8-12% of the possible SAVs in each CDK, similar to the editing rate predicted by perfect editing in the +4 to +8 window. In contrast, at a more stringent frequency cutoff of 40%, this number dropped to around ∼5% of possible SAVs (**Figure 1e**). When we performed the same analysis, but instead focused on the codon level, we found that we were able to mutate >80-90% of the amino acids in each gene at a frequency cutoff of 10%, which is higher than the expected mutation rate based on the +4 to +8 editing window (**Figure 1f**). Even at the more stringent frequency cutoff of 40%, we could mutagenize >80% of the amino acids in most genes, with the exception of *CDK7* where close to 60% of the codons were mutated (**Figure 1f**). These results suggest that, despite the limited set of transition mutations that are engineered by adenine and cytosine base editors, this base editing sensor tiling approach provides extensive mutational breadth by installing mutations at nearly every amino acid in each gene with high efficiency.

Base editors can also mutate multiple bases at once, leading to generation of compound/composite mutations where multiple amino acids are mutated *in cis* (*79*–*81*). We found that, at an editing frequency cutoff of 10%, a large fraction of the observed mutations were compound mutations (**Figure 1g**). In fact, though the number of single mutants was similar to the number expected by editing in the +4 to +8 window, the number of compound mutants exceeded the predicted value, likely as a result of the expanded editing window that we observed (**Figure 1d,g**). These results further highlight the unique ability of the sensor-based approach in providing empirical qualitative and quantitative readouts of gRNA activity, allowing for more accurate deconvolution of alleles that mediate resistance.

### Identification of known and novel resistance variants in CDK9 and CDK7

Initially focusing on the CDK9-targeting PROTACs and inhibitors, we identified CDK9 L156F as the strongest resistance variant in cells treated with the KI-CDK9d-32 PROTAC and KI-CDK9d-32N, a modified version of the PROTAC with no CRBN binding activity and thus no degradation activity **(Figure 2a-c)**. Four distinct gRNAs, each engineering the CDK9 L156F variant within the ATP-binding pocket with high efficiency, were strongly enriched in cells treated with these drugs **(Figure 2a-c)**. This variant was previously identified in a dose escalation study with BAY1251152, a different ATP-competitive CDK9 inhibitor, serving as strong orthogonal validation of our screening results (*37*, *82*). Beyond recapitulation of previously identified resistance variants, we also identified novel variants in the CDK9 ATP-binding pocket, including C106Y_E107K and D104N_C106Y—two compound variants located in the kinase linker region that interacts directly with these molecules (*62*) **(Figure 2b-c).**

**Figure 2.**
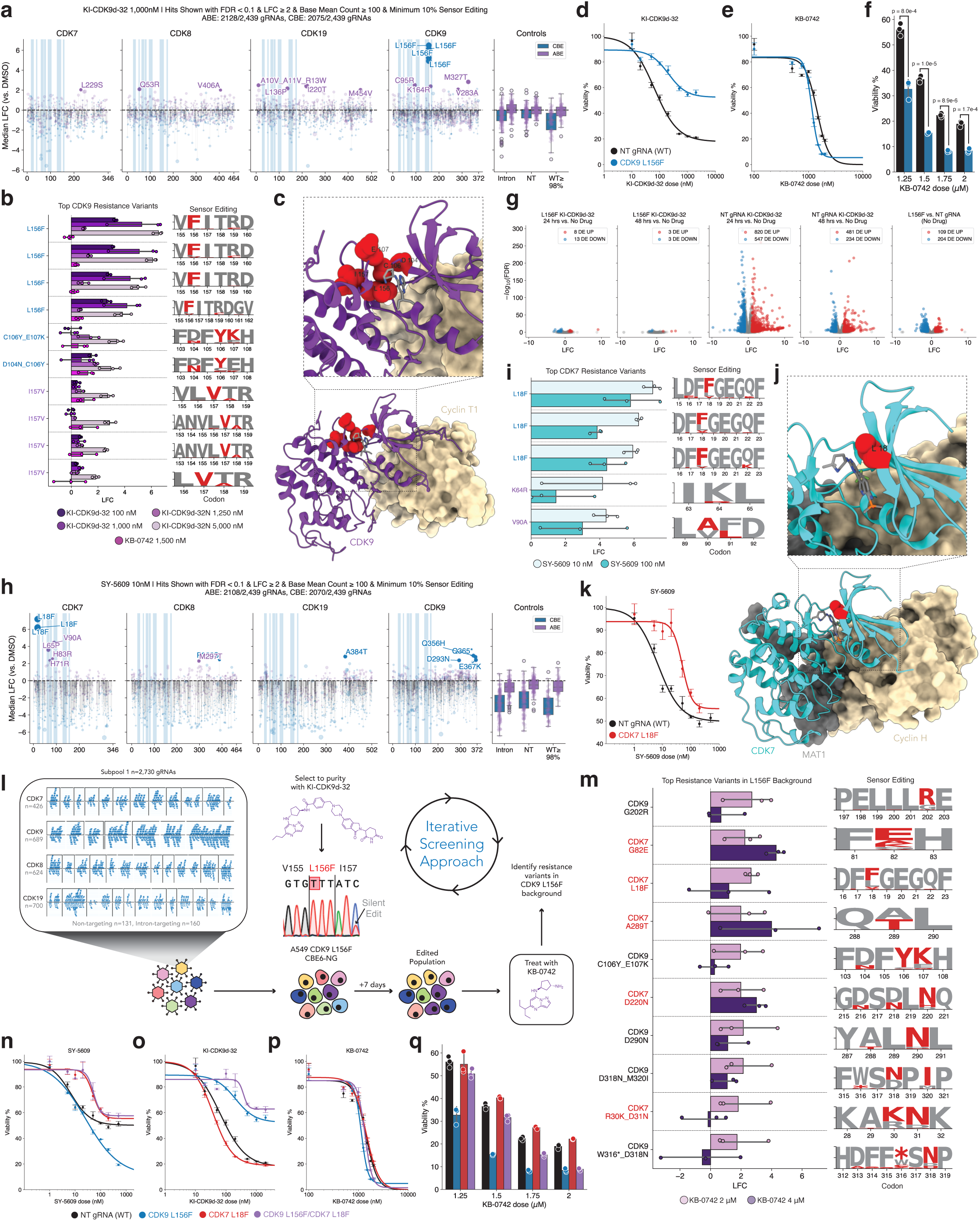
Identification and validation of resistance variants in CDK9 and CDK7. **a)** Scatterplot of median gRNA enrichment (log_2_ fold-change) in the presence of 1,000 nM KI-CDK9d-32 relative to DMSO-treated control. Each dot represents a gRNA, filtered to exclude gRNAs below 10% sensor editing and with a base mean count ≤ 100. Hits labelled with FDR < 0.1 & LFC ≥ 2. ABE screen results shown in purple; CBE in blue. Regions shaded blue indicate ATP binding site residues. **b)** Visualization of top 10 CDK9 resistance variants. Maximum likelihood estimate of sensor editing rate shown as amino acid logo plot. **c)** Structural visualization of top CDK9 resistance alleles (shown in red) in complex with KB-0742. PDB: 8K5R. **d-e)** Dose-response curve testing of CDK9 L156F and WT (non-targeting gRNA) cells in response to KI-CDK9d-32 (d) and KB-0742 (e). **f)** Quantification of dose-response curve results in (e), with p-values shown from t-test for independent means. **g)** Bulk RNA-sequencing results from CDK9 L156F and NT gRNA cells treated with KI-CDK9d-32 at designated time-points. Differentially expressed genes defined as FDR ≤ .01 and |LFC| ≥ 1. **h)** Same as (a), but showing results from treatment with low-dose SY-5609. **i)** Same as (b), but showing top 5 CDK7 resistance variants. **j)** Structural visualization of CDK7 L18 residue (red) in complex with SY-5609. PDB: 8S0T. **k)** Dose-response curve of CDK7 L18F and NT gRNA cells when treated with SY-5609. **l)** Schematic iterative screening approach. **m)** Top resistance variants from the iterative screen, as ranked by LFC. **n-p)** Dose response curves of WT, CDK9 L156F (blue), CDK7 L18F (red), and double mutant CDK9 L156F/CDK7 L18F when treated with SY-5609 (n), KI-CDK9d-32 (o), and KB-0742 (p). **q)** Barplot representation of (p).

Notably, we found that CDK9 L156F conferred resistance to both forms of the CDK9 PROTAC, KI-CDK9d-32 and KI-CDK9d-32N, but not to the ATP-competitive inhibitor KB-0742 **(Figure 2d-f)**. This is despite the high degree of structural similarity between KB-0742 and the CDK9-binding moiety of the PROTAC, as KB-0742 is an optimized version of the CDK9 binding moiety for KI-CDK9d-32 (*62*). In fact, it appeared that the CDK9 L156F mutation subtly increased the sensitivity of cells to KB-0742 **(Figure 2f)**. We posit that this difference could be due to the larger size of the PROTAC, which may enable more facile steric occlusion of the bulkier KI-CDK9d-32 from CDK9’s ATP binding pocket.

Given the central role of CDK9 in global transcriptional processes, we also performed bulk RNA-sequencing on the CDK9 L156F mutant, as well as a WT control cell line with a non-targeting (NT) gRNA, in the presence or absence of KI-CDK9d-32 **(Figure 2g)**. We found that treatment of CDK9 L156F cells with KI-CDK9d-32 induced no significant transcriptional changes. In contrast, WT cells treated with KI-CDK9d-32 show extensive transcriptional changes at both 24- and 48-hours post-treatment. Interestingly, even in the absence of the PROTAC, there are notable differences in the transcriptional profiles of CDK9 L156F cells when compared to isogenic WT cells **(Figure 2g)**.

Next, we focused on variants that were enriched in the presence of a clinically tested ATP-competitive CDK7 inhibitor, SY-5609. We identified CDK7 L18F, a variant located in a beta-sheet near the glycine-rich loop (g-loop) within the ATP binding site **(Figure 2h-k).** To our knowledge, CDK7 L18F is the first resistance variant reported for CDK7. Multiple gRNAs that engineer the CDK7 L18F variant with high efficiency were strongly enriched in the context of both low- and high-dose exposure to SY-5609 **(Figure 2h-i).** Treatment with SY-5609 led to fully homozygous, biallelic positive selection for CDK7 L18F, and subsequent dose-response testing of biallelic CDK7 L18F cells confirmed that this variant is a *bona fide* resistance variant **(Figure 2k, Supplementary Figure 3).** Other variants in CDK7 also appear to confer resistance, including V90A, located near the gatekeeper residue of the kinase (**Figure 2h-i)**.

For certain inhibitors, particularly the CDK8/19 inhibitors that we tested, no clear resistance variants emerged or were able to be validated, though we did observe enrichment of variants in both CDK8 and CDK19 in these treatment contexts **(Supplementary Figure 3)**. The high signal-to-noise ratio in our screens, the validation experiments we performed, and the orthogonal validation of previously published dose-escalation studies that confirmed our results suggest the possibility that these inhibitors are promiscuous. The polypharmacology of these compounds, which could be inhibiting multiple CDKs or other proteins at the doses we tested, would likely prevent a single point mutation from conferring resistance, as the cytotoxic effects of the drugs could still be mediated through inhibition of other proteins. As such, the base editing sensor tiling assay performed in this manner could in principle be used as a powerful method to determine the relative specificity of an inhibitor, particularly when targeting proteins with multiple structurally similar relatives.

### An iterative screening approach identifies intra-family epistatic resistance to therapy

Given the observation that CDK9 L156F confers resistance to the CDK9 PROTAC KI-CDK9d-32, but not the inhibitor KB-0742 **(Figure 2d-f)**, we hypothesized that other residues within CDK9 itself or in other CDKs would mediate sensitivity and resistance to KB-0742. To test this hypothesis, we performed an iterative screening approach to identify resistance variants against KB-0742 in the CDK9 L156F genetic background **(Figure 2l)**. This is akin to modeling the often slow emergence of a secondary resistance mutation following treatment with a second-line therapy, a process commonly observed in the presence of other targeted therapies (*35*, *36*). It is worth noting that if we had used a cDNA library of exogenous CDK mutants, we would have needed to design an entirely new library anchored on the CDK9 L156F mutation; however, the base editing sensor tiling approach allows for ‘library recycling’ by using the same library to perform this iterative screen **(Figure 2l)**.

Intriguingly, we identified several mutations in CDK7 that were enriched in CDK9 L156F cells treated with KB-0742, including the CDK7 L18F variant **(Figure 2m).** When the non-optimized lead compound for KB-0742, KI-ARv-03, was tested for kinase inhibition activity with the KinaseProfiler™ assay, CDK7/cyclin H/MAT1 was the most potently inhibited off-target, with 45% remaining activity compared to 7% remaining activity for CDK9/ cyclin T1 (*62*). In contrast, testing of the more potent KB-0742 with the HotSpot Kinase Assay found strong selectivity for CDK9 over CDK7 (*62*). Still, testing of the chemically similar lead compound KI-ARv-03 suggests some level of engagement of CDK7 by KB-0742, at least *in vitro*. Nevertheless, *in vitro* assays with purified protein may not accurately reflect the physiological environment of the cell where proteins are often involved in higher order molecular complexes that may modulate inhibitor engagement.

To formally test whether the CDK7 L18F mutation confers resistance to KB-0742 in the CDK9 L156F genetic background, we generated homozygous CDK9 L156F/CDK7 L18F double mutant cells **(Supplementary Figure 3)**. This was performed via delivery of the CDK7 L18F gRNA to CDK9 L156F cells and subsequent selection with the CDK7 inhibitor SY-5609. Dose-response testing of CDK9 L156F, CDK7 L18F, and the double mutant CDK9 L156F/CDK7 L18F with SY-5609 demonstrated that the L156F/L18F double mutant retained resistance to SY-5609 **(Figure 2n)**. Similarly, testing this collection of mutants against KI-CDK9d-32 revealed strong resistance of the double mutant, surpassing the resistance conferred by CDK9 L156F alone **(Figure 2o)**. This suggests that KI-CDK9d-32 may exert some of its cytotoxicity through degradation of CDK7. Finally, testing of this collection of mutants against KB-0742 indicated that none of these mutants provide treatment resistance **(Figure 2p-q).** However, the additional sensitivity of CDK9 L156F cells to KB-0742 seems to be brought back to baseline levels in the double mutant **(Figure 2p-q)**. Altogether, these results may suggest that either (1) KB-0742 is still able to engage and inhibit CDK9 and/or CDK7 in the presence of these mutations, or (2) at the doses needed to achieve cytotoxicity, KB-0742 engages multiple targets, and ablation of CDK9 and CDK7 binding is insufficient to provide resistance to this compound.

### *In vivo* validation of resistance variants in CDK9 and CDK7

Next, we sought to determine whether the CDK9 L156F and CDK7 L18F variants confer resistance in a physiologically relevant, *in vivo* environment. Due to the near perfect conservation of *CDK9* and *CDK7* between human and mouse, we were able to engineer these mutations in murine B-cell acute lymphoblastic leukemia (B-ALL) cells driven by the BCR-ABL oncogene in a homozygous *Arf* null background (*83*). We engineered these variants by transient electroporation of mRNA encoding CBE6-NG into B-ALL cells harboring CDK9 L156F or CDK7 L18F gRNAs **(Figure 3a)**. Importantly, this system allowed us to test the response of WT or CDK-mutant B-ALL cells to KI-CDK9d-32 and SY-5609 in a syngeneic, immunocompetent host **(Figure 3a).** We observed robust positive selection for both CDK9 L156F and CDK7 L18F in cells treated with KI-CDK9d-32 and SY-5609, respectively, demonstrating that these variants confer resistance in both a different cell type and species context **(Figure 3b-c)**.

**Figure 3.**
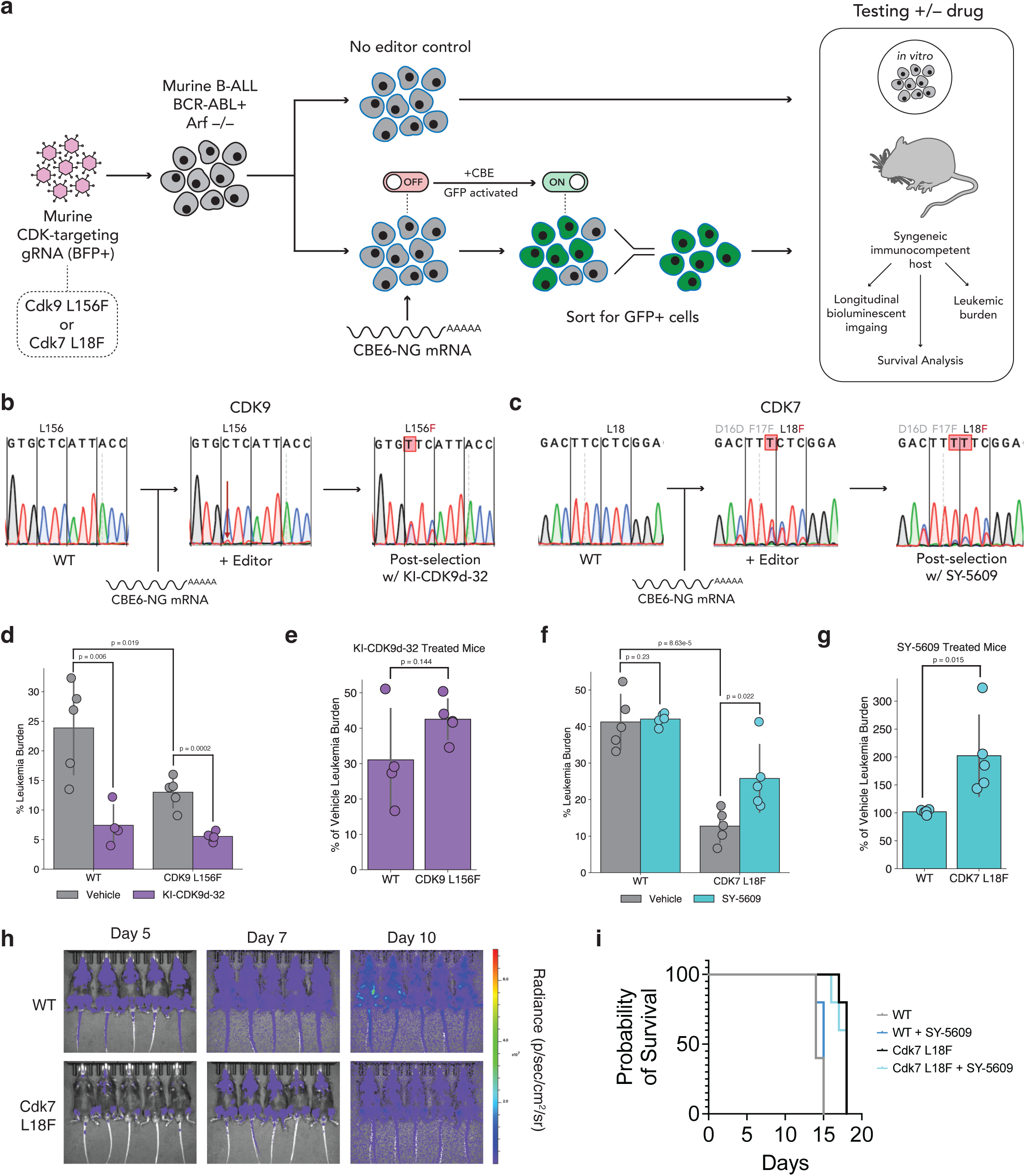
*In vivo* validation of resistance to CDK-targeting therapies. **a)** Schematic of *in vivo* testing strategy. Murine B-ALL cells are transduced with gRNA lentivirus, before electroporation, or lack thereof, with CBE6-NG mRNA and sorting for edited cells. Mice are injected with WT and mutant cells and leukemic burden is monitored over time. **b)** Sanger sequencing analysis of Cdk9 L156F in mouse cells before editor electroporation, after electroporation, and after selection with KI-CDK9d-32. **c)** Sanger sequencing analysis of Cdk7 L18F in mouse cells before editor electroporation, after electroporation, and after selection with SY-5609. Quantification of leukemic burden in the spleen as assessed via flow cytometry at day 10 post-injection with cells. P-values indicate t-test for independent means. **e)** Leukemic burden for KI-CDK9d-32-treated mice quantified as a percentage of the leukemic burden of vehicle-treated control mice. **f)** Same as (d), but for Cdk7 L18F mutant lines and matched WT control, treated with vehicle or SY-5609. **g)** Same as (e), but for SY-5609-treated mice. **h)** Longitudinal bioluminescent imaging of mice injected with WT or Cdk7 L18F B-ALL cells in the vehicle treatment condition. Images at day 5, 7, and 10 post-injection. **i)** Survival curves of mice injected WT or Cdk7 L18F B-ALL cells and treated with vehicle or SY-5609; n=5 mice per condition.

We first tested the leukemic progression of WT and CDK9 L156F B-ALL cells in the presence or absence of KI-CDK9d-32. It is worth noting that KI-CDK9d-32 had not been previously tested *in vivo*. We measured the leukemic burden of mice using flow cytometry by quantifying fluorophore-positive blasts in the spleens of mice from the different cohorts. In both genotypes, we observed a significantly lower leukemic burden in KI-CDK9d-32-treated animals compared to vehicle-treated controls **(Figure 3d)**. However, we also observed a significantly lower leukemic burden in mice injected with CDK9 L156F cells compared to those injected with WT B-ALL cells **(Figure 3d)**. This suggests that the CDK9 L156F variant could produce a fitness defect in these cells that slows leukemic progression. In agreement with this observation, previous work has shown that the CDK9 L156F variant results in reduced CDK9 kinase activity (*37*). When we normalized the leukemic burden of KI-CDK9d-32-treated animals to that of vehicle-treated controls, we did observe increased burden in the CDK9-mutant setting, indicating that the CDK9 L156F does provide some degree of resistance to KI-CDK9d-32 in this setting **(Figure 3e)**. However, survival analysis was not possible with this cohort due to the toxicity of KI-CDK9d-32 treatment.

Next, we tested whether the CDK7 L18F variant provides resistance to SY-5609 in the same B-ALL system. Similar to CDK9 L156F, we found a nearly four-fold lower leukemic burden in mice injected with CDK7 L18F cells compared to those injected with WT B-ALL cells, reflecting a potential fitness defect in the CDK-mutant line **(Figure 3f)**. Notably, when comparing the leukemic burden within a given genotype (WT or CDK7 mutant), we observed significantly higher leukemic burden in the CDK7-mutant line when animals were treated with SY-5609, suggesting a resistance phenotype *in vivo* despite the fitness cost conferred by this mutation **(Figure 3f-g).** This could be the result of growth suppression of non-leukemic cells in the presence of SY-5609. Moreover, longitudinal bioluminescent imaging and survival analyses further support a fitness defect in the CDK7 mutant line **(Figure 3h-i).** Altogether, these results demonstrate an *in vivo* resistance phenotype for CDK7 L18F, as well as a fitness defect in this cellular context.

### Identifying general and therapeutic modality-specific CDK12/13 resistance variants

Given our success in identifying known and novel resistance variants in CDK7 and CDK9, we expanded our approach to identify resistance variants in other CDKs. We started by extending our investigation to subpool 2, which contains CDK12 and CDK13, two paralogs involved in transcriptional elongation and termination **(Figure 1b)**. After mutagenizing CDK12/13, we treated cells with 3 molecules with different mechanisms of action: (1) CDK12-IN-2 (*71*), an ATP-competitive CDK12/13 inhibitor, (2) BSJ-4-116 (*39*), a CDK12/13 PROTAC, and (3) HQ461 (*38*), a molecular glue degrader that facilitates an interaction between CDK12/13 and DDB1 to mediate the ubiquitination of cyclin K.

Once again, as an orthogonal validation of the efficacy of our approach in identifying *bona fide* resistance variants, we observed strong concordance between the resistance variants identified through dose escalation studies and those that we identified with base editing. First, we identified CDK12 I733V as the most enriched variant when cells were treated with BSJ-4-116 **(Figure 4a)**. A dose escalation study independently identified CDK12 I733V as a resistance variant against this PROTAC (*39*). Second, we found that CDK12 G731K was the most enriched variant in the CBE arm of the screen, and the second most enriched variant overall, when cells were treated with HQ461 **(Figure 4b-c)**. This finding also corresponds directly with the findings of an independent dose escalation study that identified two mutations in the same CDK12 amino acid (G731E and G731R) that confer resistance to HQ461 (*38*). Notably, other studies have identified DDB1 and CUL4B mutations that confer resistance to other CDK12 molecular glue degraders (*84*).

**Figure 4.**
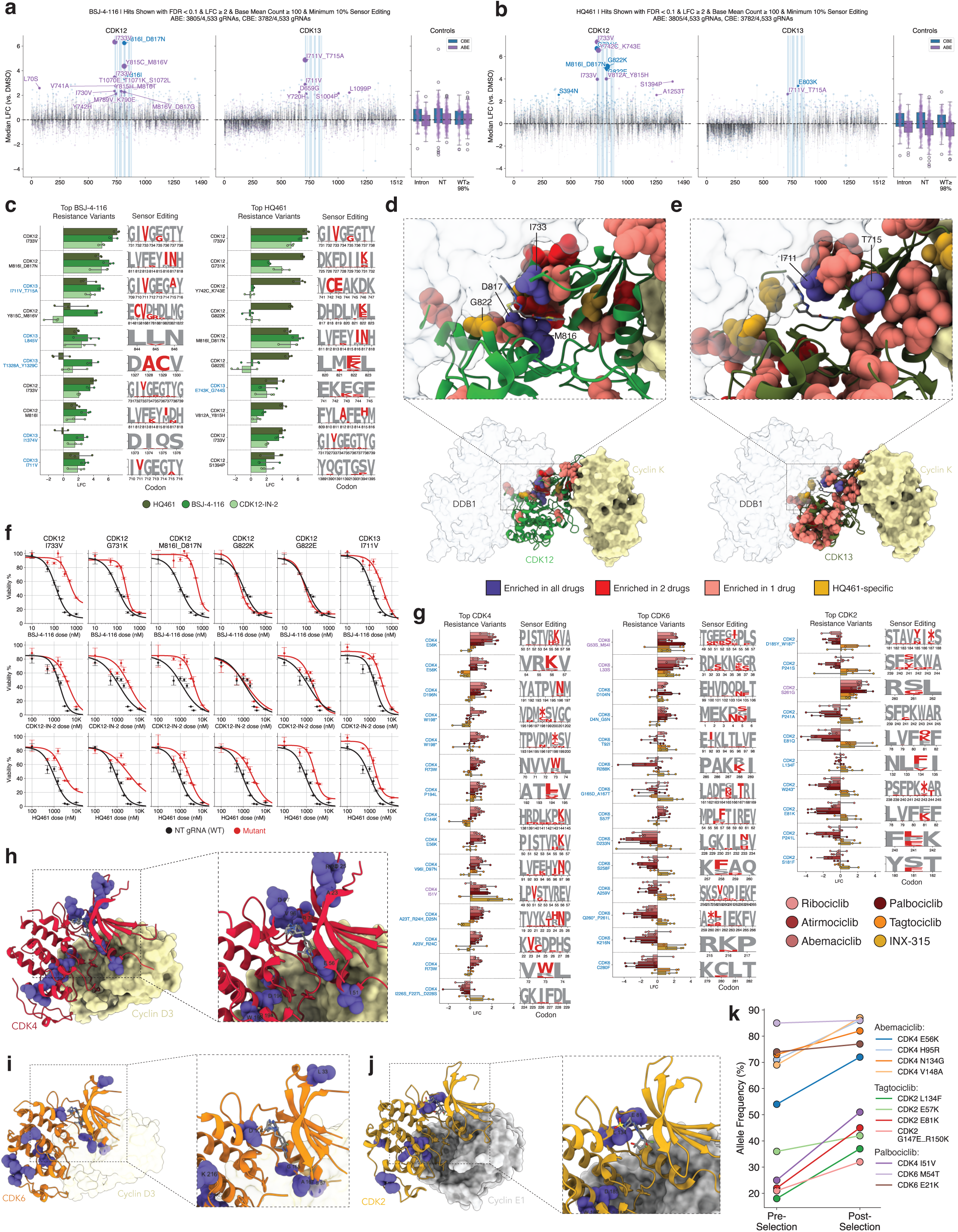
CDK-drug interaction & resistance mapping for CDK12/13 and CDK2/4/6. **a)** Scatterplot of median gRNA enrichment (log_2_ fold-change) in the presence of BSJ-4-116 relative to DMSO-treated control. Each dot represents a gRNA, filtered to exclude gRNAs below 10% sensor editing and with a base mean count ≤ 100. Hits labelled with FDR < 0.1 & LFC ≥ 2. ABE screen results shown in purple; CBE in blue. Regions shaded blue indicate ATP binding site residues. **b)** Same as (a), but for HQ461-treated cells. **c)** Visualization of the top resistance variants in BSJ-4-116- and HQ461-treated cell pools. Maximum likelihood estimate of sensor editing rate shown as amino acid logo plot. **d)** Visualization of enriched residues in CDK12 in complex with HQ461 and DDB1. Colors indicate enrichment in 1 (salmon), 2 (red), or all drugs (purple), or HQ461 specifically (gold). PDB: 8BUG. **e)** Same as (d), but visualizing CDK13 hits. PDB: 7NXJ (with DDB1 superimposed from PDB: 8BUG). **f)** Dose-response curves of indicated variants when treated with BSJ-4-116 (top), CDK12-IN-2 (middle), and HQ461 (bottom). **g)** Visualization of top resistance variants in CDK2/4/6. Showing variants with FDR < .01 in at least one treatment condition. Maximum likelihood estimate of sensor editing rate shown as amino acid logo plot. **h-j)** Structural visualization of CDK4-cyclin D3 bound to abemaciclib (h), CDK6-cyclin D3 bound to abemaciclib (i), and CDK2-cyclin E complex bound to PF-06873600 (j), with resistance variants highlighted in purple. PDB: 7SJ3, 5L2S, 7KJS. Cyclin D3 in CDK6 structure (i) is superimposed from CDK4 structure (7SJ3). **k)** Endogenous allele frequency of CDK4, CDK2, and CDK6 variants before and after selection with 5 µM Abemaciclib, 10 µM Tagtociclib, or 10 µM Palbociclib. Sanger sequencing quantified with EditR software.

In addition, we identified novel resistance variants in both CDK12 and CDK13 in the context of all three mechanistically-distinct small molecules we tested, with a particular enrichment for variants near the g-loop, hinge, and linker region in the ATP-binding cleft **(Figure 4c-e, Supplementary Figure 4)**. CDK12 and CDK13 are paralogs with a high degree of sequence and structural conservation, leading us to hypothesize that these molecules may bind to both paralogs. Supporting our hypothesis, we found that resistance variants can emerge in either protein, potentially suggesting a compensatory mechanism—if one paralog is able to avoid inhibition or degradation, it may functionally compensate for the inhibition of the other paralog.

Notably, we also identified variants that promote resistance to mechanistically-distinct CDK12/CDK13 inhibitors, including CDK12 G822K, which specifically conferred resistance to the HQ461 molecular glue degrader but not to CDK12-IN-2 or BSJ-4-116 **(Figure 4c)**. These variants are located at the DDB1-CDK12 interface (*85*), suggesting that resistance is achieved through disruption of this interface, which would prevent cyclin K degradation, rather than through disruption of drug binding **(Figure 4d-e)**.

Next, we performed individual validation experiments for 6 of these resistance variants, including 5 CDK12 variants (I733V, G731K, M816I_D817N, G822K, G822E) and 1 CDK13 variant (I711V), by generating near-homozygous variant cell lines via selection with HQ461 **(Figure 4f, Supplementary Figure 4).** In dose-response curves, each of these variants conferred significant resistance to all 3 of the tested drugs, with the exception of CDK12 G822E and G822K, which only conferred resistance to HQ461, confirming the unique resistance mechanism of these variants **(Figure 4f)**. Of note, both CDK12 I733V and CDK13 I711V align to the same structural position in beta-sheet I prior to the g-loop of CDK12/13, suggesting a convergent mechanism of resistance in both paralogs. We confirmed that the CDK13 I711V gRNA does not produce off-target editing in CDK12 and that this resistance phenotype is in fact mediated through the mutation in CDK13 **(Supplementary Figure 4).**

### A CDK-drug interaction map for CDK2/4/6

Next, we extended our analysis to CDK2/4/6, the cell-cycle CDKs included in subpool 3 **(Figure 1b)**. We mutagenized these CDKs and then sought to identify resistance variants against multiple clinically approved, standard-of-care CDK4/6 inhibitors, as well as inhibitors of CDK4/6 and CDK2 currently in clinical trials. This includes Palbociclib, Abemaciclib, and Ribociclib, clinically approved CDK4/6 inhibitors commonly used in treating breast cancers, as well as Atirmociclib, a novel CDK4-selective inhibitor, and two CDK2 inhibitors in clinical trials, Tagtociclib and INX-315. While several primary and secondary mechanisms of resistance have been previously identified (*26*, *40*–*42*), the molecular underpinnings of cancer resistance and progression to CDK4/6i remains to be elucidated in about half of the cases. As such, we sought to identify variants in CDK2/4/6 that could confer resistance to these therapies.

We observed similar resistance profiles for the three clinically approved CDK4/6 inhibitors, with a number of hits in both CDK4 and CDK6 **(Figure 4g, Supplementary Figure 5).** The majority of significantly enriched variants localized to the ATP-binding pocket, though we also observed hits in multiple areas of the C-lobe of CDK4/6 that are distal to this pocket **(Figure 4h-i)**. Of particular note, two gRNAs that engineer the CDK4 E56K variant with high efficiency were enriched in the presence of all three clinically approved CDK4/6 inhibitors, as well as in the presence of the CDK4-selective inhibitor, Atirmociclib (**Figure 4g-i)**. This variant is located in the αC-helix of CDK4, which is the structural motif involved in cyclin-binding, suggesting potential disruption of the CDK4-cyclin D interaction, or alterations to the structure of the CDK-cyclin complex that prevent inhibitors from binding. Given the importance of the CDK4/6-Cyclin D-p27 interaction in modulating the activity of CDK4/6, where p27 acts as a CDK inhibitor, these variants could modulate this complex to generate the resistance phenotype (*86*)

Similarly, we identified multiple resistance variants in CDK2 that arise in response to treatment with INX-315 or Tagtociclib. This includes CDK2 E81Q, located in the hinge region of the ATP-binding cleft, and CDK2 L134F, which notably aligns directly with CDK9 L156F **(Figure 4g, j)**.

In addition, we performed arrayed validation of the resistance phenotype of a selection of variants in or near the ATP binding pockets of CDK4 (E56K, H95R, N134G, V148A, I51V), CDK2 (L134F, E57K, E81K, G147E_R150K), and CDK6 (M54T, E21K). We found a significant increase in the endogenous allele frequency of these variants after selection with Abemaciclib, Tagtociclib, or Palbociclib, excluding the CDK6 alleles whose frequency remained relatively consistent, confirming the selective advantage of these variants in the presence of these inhibitors **(Figure 4k)**.

### CDK residue essentiality revealed by multiplexed base editing

Our multiplexed mutagenesis approach also provides the ability to quantify residue essentiality in the absence of perturbations like drug selection **(Figure 5a)**. By comparing the frequency of CDK mutants at the “T0” timepoint relative to the plasmid library, we can map the fitness effects of mutations in each of the CDKs **(Figure 5a)**. To do so, we calculated the average value of the essentiality score (log_2_ fold-change of the T0 v. plasmid library) at each mutated amino acid position to generate residue-level scores for each CDK. We observed a moderate correlation (R=0.66) when comparing the average residue essentiality scores between the ABE and CBE screens, likely reflecting the differing phenotypic effects of the mutations generated by each editor, as well as technical noise **(Figure 5b)**.

**Figure 5.**
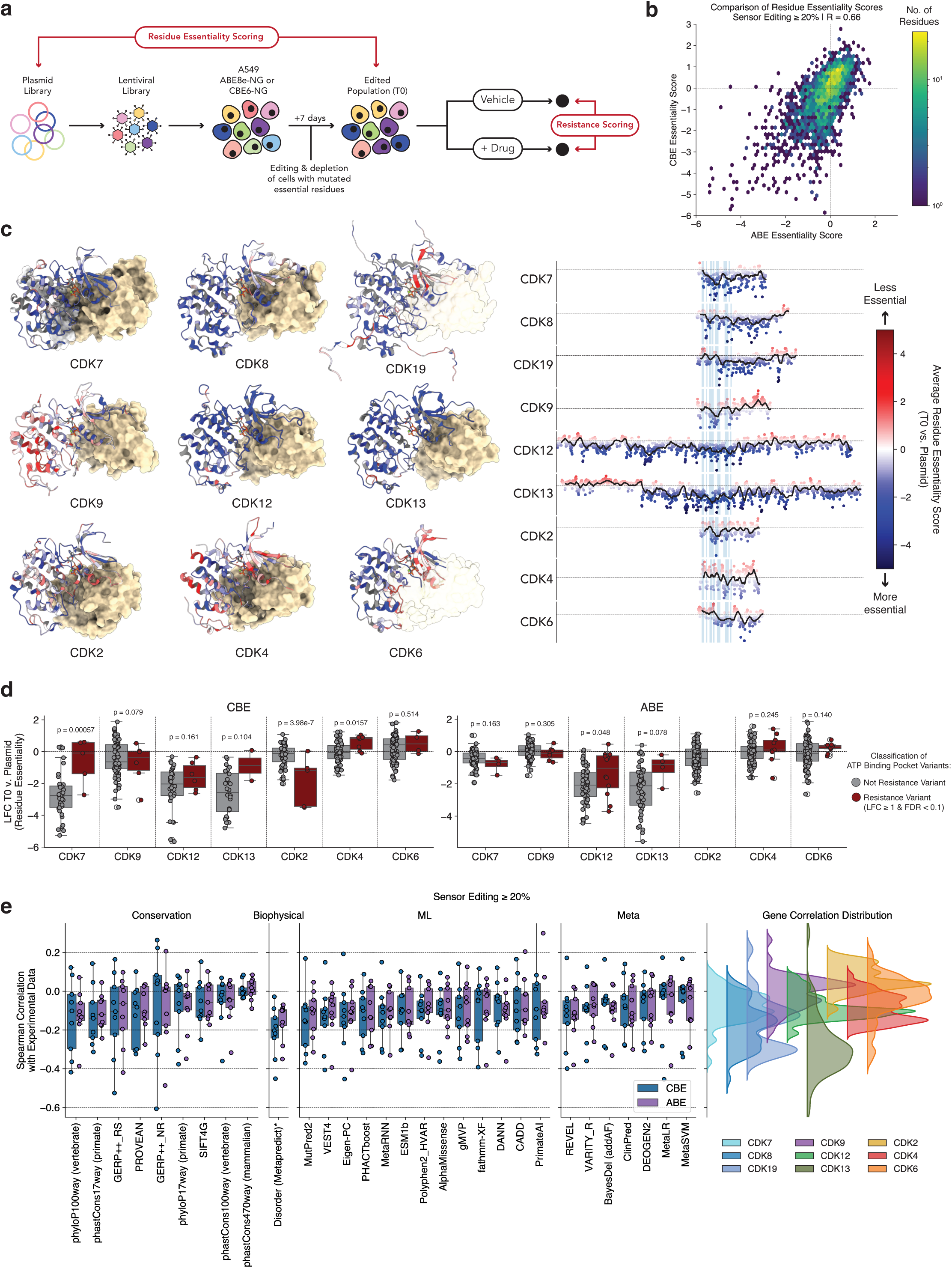
CDK residue essentiality revealed by multiplexed base editing. **a)** Schematic of the comparison used to generate residue and mutation essentiality scores via comparison of the frequency of mutants at the “T0” timepoint with the Plasmid Library. **b)** Binned scatterplot showing the correlation between residue essentiality scores from ABE and CBE screens. Pearson correlation (R) = 0.66. **c)** Right: residue essentiality scores, averaged across gRNAs targeting a given residue with ≥20% editing efficiency, plotted across the protein sequence of each CDK. Shaded blue regions indicate ATP binding domains; proteins are aligned such that the start site of each ATP binding domain is consistent. The dashed black line indicates a residue essentiality of 0, while the shaded black line shows a 20 amino acid moving average. Left: structural visualization of the residue essentiality scores for each CDK. Colors are normalized so that the minimum value (blue) is -1 and the maximum value (red) is 1. PDB/Alphafold codes: CDK7, 8S0T; CDK8, 5CEI; CDK19, Q9BWU1 (Alphafold); CDK9, 8K5R; CDK12, 8BUG; CDK13, 7NXJ; CDK2, 1QMZ; CDK4, 7SJ3; CDK6, 5L2S. CDK structures are aligned and depicted with ATP from 1QMZ. Transparent cyclin structures indicate non-presence in the original structure. **d)** Boxplot of residue essentiality for resistance variants (red) and non-resistance variants (grey). P-values derived from t-test for independent means. **e)** Boxplots showing the spearman correlation between variant effect predictor (VEP) score and mutation fitness score for gRNAs engineering mutation with at least 20% editing efficiency. Right, kernel density histogram of spearman correlation values for each gene across the tested VEPs.

Next, we used the resulting data to build high-resolution structure-function maps of each CDK **(Figure 5c)**. As expected, we consistently observed depletion of the structured ATP binding domains in each kinase, indicating that many of these residues play essential roles in kinase function. This effect was particularly pronounced in the larger CDKs, CDK12 and CDK13, which contain long unstructured N- and C-terminal tails. In these unstructured regions, we observed significantly less residue essentiality when compared with the more structured kinase domains **(Figure 5c)**.

However, mutations in the ATP binding domain did not deplete uniformly **(Figure 5c)**. Given that many of these kinases are essential for cell function, we hypothesized that resistance variants must balance ablation of inhibitor binding with the maintenance of sufficient kinase activity to allow for cell survival. To test this, we analyzed patterns of residue essentiality in the ATP binding domain of each CDK, quantifying the difference between resistant and non-resistant variants **(Figure 5d)**. With the exception of CDK2, we found that resistance variants in the ATP binding cleft tend to occur in residues that are more readily able to tolerate mutations. This effect is particularly pronounced in the CDK7 resistance variants identified in the CBE arm of the screen (including CDK7 L18F), as well as in the CDK12/13 resistance variants **(Figure 5d)**.

More generally, we assessed whether any existing variant effect prediction (VEP) tools could explain the patterns of essentiality that we observe in our data. These tools provide a pathogenicity score, with higher scores indicating that a mutation has a more damaging effect. Based on this premise, we would expect the predicted pathogenicity scores to be anticorrelated with mutant fitness, with lower scores indicative of a more damaging mutation. Indeed, we observed a modest negative correlation when we compared our experimental data with these predictions **(Figure 5e)**. Nevertheless, we observed a markedly high degree of variance in the fidelity of these predictions on a gene by gene basis, indicating that these VEP tools are unable to accurately predict and recapitulate empirically determined mutation fitness scores at scale. One possible explanation for the lack of predictive power of VEPs in this context could be a failure to adequately consider protein-protein interaction interfaces, which may not be fully captured by these models. We note that in this particular dataset, the best predictor of mutation fitness is in fact the simple biophysical prediction of amino acid disorder **(Figure 5e)**.

### Pan-CDK patterns of resistance to CDK inhibitors, PROTACs, & molecular glue degraders

Beyond delineating a catalog of resistance variants, we investigated whether we could identify common patterns of resistance across the CDKs that we profiled. We reasoned that systematic mutagenesis of a family of closely related proteins would uniquely enable us to align and compare the resistance profiles of the CDKs. To identify shared and divergent patterns of resistance across the CDK family, as well as other kinases, we performed both structural and sequence-based alignments.

First, we performed a structural alignment using the KLIFS universal kinase structural alignment indexing system, which provides a method of directly comparing the ATP-binding pockets of all of the kinases in the human kinome (*87*, *88*). Initially, we compared the observed resistance spectra in the CDKs with clinically-observed resistance variants in commonly targeted kinases in oncology via this structural alignment **(Figure 6a-b)**. To do so, we queried the OncoKB database, identifying 42 unique mutations across 8 kinases that were located in the ATP-binding pocket as defined by KLIFS **(Figure 6c)**. We found that these resistance mutations were predominantly localized to the Gatekeeper, Hinge, and Linker regions of these kinases, with some resistance mutations in g-loop and other regions **(Figure 6d).** Consistent with clinical data, we also observed an enrichment of resistance mutations in the Hinge and Linker region **(Figure 6e)**. However, we also identified other common sites of resistance, including in residues within the g-loop region, the beta sheets prior to the xDFG motif, and the αC-helix **(Figure 6e)**. Importantly, commonly observed resistance hotspots were not due to editing efficiency biases, as we observed a relatively homogeneous editing rate of amino acids in the ATP binding site, as defined by KLIFS.

**Figure 6.**
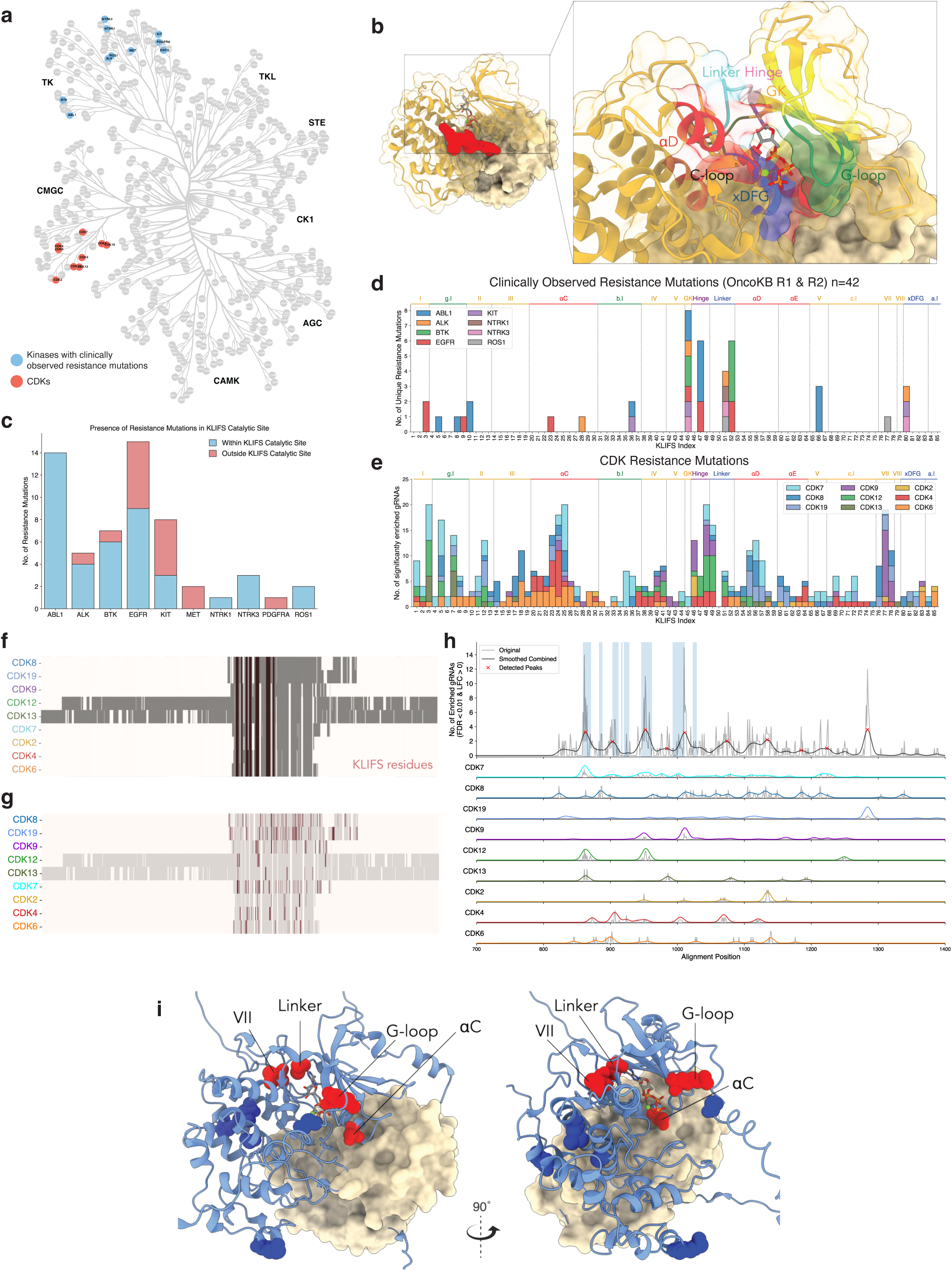
Pan-CDK patterns of therapy resistance. **a)** Tree visualization of the human kinome, with the profiled CDK family members and the OncoKB kinases with clinical resistance information highlighted in red. **b)** Visualization of the KLIFS classification of the ATP binding pocket shown on peptide-bound (red), phosphorylated CDK2 structure. PDB: 1QMZ. **c)** Quantification of the presence of OncoKB resistance mutations within or outside of the KLIFS-defined kinase catalytic site. **d)** Stacked barplot of the number of unique OncoKB resistance mutations at each KLIFS site. **e)** Barplot of the number of significantly enriched CDK-targeting gRNAs (FDR < .01) at each KLIFS site within the ATP-binding site. Colors indicate gene. **f)** Visualization of multiple sequence alignment among the profiled CDKs, with the KLIFS-defined catalytic site highlighted in red. **g)** Same as (f), but showing the significantly enriched gRNAs in red. **h)** Quantification of the number of enriched gRNAs at each aligned position in the multiple sequence alignment (for alignment positions 700 to 1400). Curves indicate kernel density estimate; red “x” marks indicate peaks detected by the peak-finding algorithm. **i)** Visualization of peaks detected in (h), with residues within the KLIFS catalytic motif highlighted in red, and residues outside highlighted in blue. Alphafold structure of CDK19 (Q9BWU1) with cyclin superimposed.

Owing to the high degree of sequence conservation among the CDKs, we also performed a multiple sequence alignment of the CDKs that we profiled to identify aligned motifs that commonly promote resistance to CDK inhibitors **(Figure 6f-g)**. In concordance with the KLIFS-based structural alignment, we observed multiple distinct peaks of enrichment in the ATP-binding pocket, including in the g-loop, beta sheet VII, the Linker region, and the αC-helix **(Figure 6h-i)**. Notably, we also observed the presence of shared enrichment outside of the ATP-binding pocket, particularly in the C-terminal regions of these CDKs, suggesting potentially allosteric sites or mechanisms of resistance. Despite patterns of similarity among the CDKs that we profiled, there does appear to be extensive heterogeneity in the resistance spectra of the CDKs, likely reflecting the divergence of the CDK family members.

### Clinical evidence of impactful CDK mutations in breast cancer patients

CDK4/6 inhibitors, in combination with aromatase inhibitors, represent the current standard of care regimen in the frontline metastatic setting and for high-risk, early stage hormone receptor positive breast cancers (*15*–*18*, *21*, *22*, *89*). Accordingly, given our preliminary results showcasing specific CDK mutagenic patterns to shape therapeutic sensitivity to CDK4/6i, we sought to map our findings in patients affected by HR+/HER2-breast cancer, and determine whether the behavior of these mutations in base editing sensor tiling screens was correlated with clinical outcomes in those treated with CDK4/6 inhibitors.

We interrogated a large clinical cohort of patients with breast cancers which received tumor sequencing using the FDA-cleared, tumor-normal, MSK-IMPACT assay (*40*, *90*, *91*), which provides sequencing coverage spanning the CDK4 and CDK6 exonic regions. From this patient subset, we selected two patients that were treated with the CDK4/6 inhibitor palbociclib **(Figure 7a-b)**. These two patients, both with alterations in CDK6, experienced markedly different responses to palbociclib treatment. The first patient, affected by HR+/HER2-breast cancer, suffered from a metastatic tumor recurrence on adjuvant endocrine therapy, for which she received a tumor sequencing revealing a clonal CDK6 R46Q variant, together with a mono-allelic *FAT1* mutation without annotated functional significance (variance of unknown significance). Following front-line treatment with palbociclib with anastrozole, she showcased a rapid tumor progression after 7.8 months of treatment **(Figure 7a)**. In our base editing sensor tiling screens, we observed significant enrichment in the presence of palbociclib for three gRNAs with editing activity at the R46 codon, suggesting that mutations at this codon are resistance alleles, in agreement with the observed clinical resistance **(Figure 7c).** Notably, this patient also had an alteration in *FAT1*, a gene whose loss has been shown to elevate CDK6 levels and contribute to resistance to CDK4/6 inhibition (*26*). This raises the interesting possibility of cooperation between *FAT1* alterations and *CDK6* mutations contributing to resistance to CDK4/6 inhibitors by, for example, elevating mutant CDK6 levels to compensate for impaired kinase activity.

**Figure 7.**
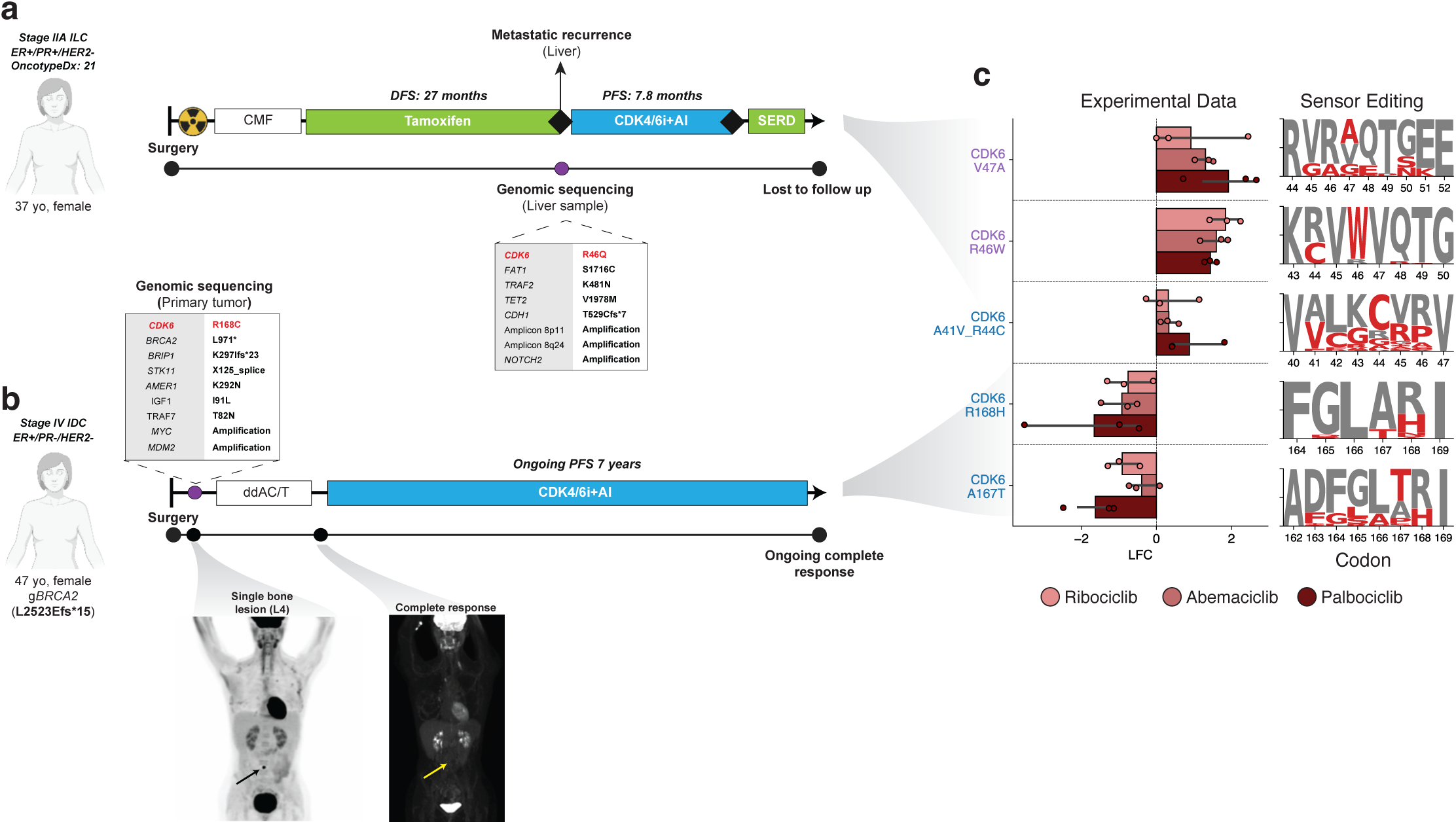
Clinical evidence of impactful CDK mutations in breast cancer patients. **a-b)** Clinical vignettes of breast cancer patients with CDK6 mutations. Patient (a) experienced relapse on CDK4/6i, while patient (b) has ongoing PFS with CDK4/6i. ILC, invasive lobular carcinoma; IDC, invasive ductal carcinoma; CMF, cyclophosphamide, methotrexate, and fluorouracil; DFS, disease-free survival; PFS, progression-free survival; AI, aromatase inhibitor; CDK4/6i, CDK4/6 inhibitor (palbociclib); SERD, selective estrogen receptor degrader; ddAC/T, Dose-dense doxorubicin and cyclophosphamide with paclitaxel. **c)** Screening data for significantly enriched or depleted gRNAs (FDR < 0.1 in palbociclib vs. DMSO) with editing activity at CDK6 R46, or CDK6 R168. Maximum likelihood estimate of sensor editing rate shown as amino acid logo plot. Blue label, CBE screening data; Purple label, ABE screening data.

In contrast, the second patient, a germline *BRCA2* mutated carrier affected by oligo-metastatic breast cancer, received primary breast surgery, and adjuvant chemotherapy followed by palbociclib together with aromatase inhibitors. Tumor sequencing performed on a metastatic bone lesion revealed a clonal CDK6 R168C mutation. Notably, she experienced a long-term benefit, with an ongoing complete response after 7 years of treatment **(Figure 7b)**. Similarly, in our base editing sensor tiling screens, we observed significant depletion of in the presence of palbociclib for two gRNAs with editing activity at the R168 codon, suggesting that mutations at this codon could confer increased sensitivity to the drug in agreement with the strong clinical response of this patient **(Figure 7c).**

Though these clinical vignettes represent only two patients, they suggest that mutations in the CDKs do occur in cancer patients and could contribute to the response to therapies targeting these proteins. As additional CDK inhibitors progress to the clinic, and as clinical sequencing efforts expand, a fuller picture of resistance mediated through mutation of these target proteins could emerge. Moreover, the present work could contribute to delineating the genomic features shaping therapeutic sensitivity to CDK4/6i, which has a relevant clinical importance for personalizing escalated treatment regimens for resistance variants.

## Discussion

In this study, we established a generalizable framework that integrates precision genome editing, mechanistically diverse therapeutics, and computational sequence-structure-function analysis to map protein essentiality and potential druggability at single amino acid resolution. We applied this framework in human cancer cells to systematically map residue essentially and the resistance landscape of 9 members of the CDK family against 15 mechanistically distinct targeted cancer therapies. By treating mutagenized cells with ATP-competitive inhibitors, PROTACs, and molecular glue degraders, we identified dozens of resistance-conferring variants, recapitulating known resistance alleles while uncovering a large repertoire of novel mutations. This includes the identification of novel resistance variants in clinically relevant targets, including CDK9, CDK7, CDK12/13, and CDK2/4/6. Moreover, by leveraging the base editing sensor architecture (*43*, *56*) to couple gRNA behavior with editing activity, we obtained new insights into endogenous residue essentiality and accurately resolved residue- and mechanism-specific differences in the resistance spectra among agents targeting the same protein, all at single amino acid resolution. These results show that base editing sensor tiling screens can serve as a generalizable, high-resolution strategy to predict and validate resistance mechanisms before they potentially emerge in the clinic.

Our findings highlight several conceptual advances. First, by performing mutagenesis across multiple CDK family members in parallel, we uncovered recurrent resistance motifs in regions that are commonly mutated in other resistant kinases, such as the Hinge and Linker regions, as well as less frequently observed resistance sites, including the g-loop region, beta sheet VII, the αC-helix, as well as regions distal to the ATP binding cleft. This cross-family view underscores the potential for comparative mutagenesis to not only inform on-target resistance, but also to reveal shared vulnerabilities exploitable in drug design in a more generalizable manner.

Second, our iterative screening strategy—an approach made uniquely facile by base editing—modeled the clinical reality of compound resistance, where tumors accumulate sequential mutations under therapeutic pressure. By screening mutants in a CDK9 L156F genetic background, we identified intra- and intergenic epistatic interactions, including selection for multiple CDK7 mutations. We note that this result is in concordance with previous testing indicating CDK7 as a potential off-target for KB-0742, demonstrating that mutagenesis of related family members in parallel can help reveal inadvertent polypharmacology.

Third, we validated key resistance alleles in an immunocompetent *in vivo* model, confirming that variants such as CDK9 L156F and CDK7 L18F confer resistance across cellular and organismal contexts. Importantly, we observed that resistance alleles can impose fitness trade-offs, with CDK9 L156F reducing leukemic progression despite conferring partial drug resistance. This balance between drug evasion and catalytic kinase impairment suggests that resistance mutations may not simply restore kinase function, but rather reconfigure signaling outputs in ways that shape tumor evolution.

Fourth, we identify multiple mutations in CDK2/4/6 that confer resistance or sensitivity to CDK4/6 and CDK2 inhibition. These findings are clinically relevant in the context of breast cancer therapy, supported by the preliminary identification of mutations in patients whose clinical outcomes correlate with the experimental behavior of these variants.

Our study also reveals modality-specific resistance signatures. For example, resistance to the molecular glue degrader HQ461 included residues that clustered at protein–protein interaction interfaces, distinct from the ATP-binding site mutations enriched under kinase inhibitor or degrader pressure. This finding emphasizes that resistance landscapes are shaped as much by drug mechanism as by target structure, and that drug modality must be considered when anticipating evolutionary trajectories of resistance.

Despite its advantages, base editing sensor tiling mutagenesis has clear limitations. The mutational depth achievable is largely constrained to transition mutations, leaving gaps in coverage compared with saturation mutagenesis approaches. At the same time, the breadth of mutagenesis across nearly every codon, the relative homogeneity and high efficiency of editing, and the ability to capture compound mutations represent strengths unique to this approach.

Another limitation is that our work was conducted primarily in A549 human lung adenocarcinoma cells. While we observed strong concordance with dose-escalation studies and extended our main findings to a murine syngeneic leukemia model, therapy response and resistance patterns may be shaped by lineage- or context-specific dependencies. Though we find that A549 cells are sensitive to all of the drugs we tested, selecting specific cell lines where each CDK is a particular dependency would allow for better calibration of the doses necessary to achieve selection, potentially improving the signal-to-noise ratio in identifying resistance variants. Future work extending these approaches to genetically-engineered mouse models that allow for genetic or pharmacological inducible and reversible inhibition of specific CDKs (*92*, *93*), patient-derived models, or more genetically and phenotypically diverse *in vivo* tumor systems will be important for translating variant maps into clinical predictions. Moreover, mutation of the target of inhibition represents only a single mechanism of resistance. Coupling these mutagenesis approaches with cancer-focused or genome-wide screens and other methods would provide a more complete picture of the diverse molecular paths that cells can take to achieve resistance.

Taken together, these findings establish a generalizable framework that integrates precision genome editing, mechanistically diverse therapeutics, and computational sequence-structure-function analysis that can be used for anticipatory resistance mapping in oncology. By prospectively identifying resistance alleles across a therapeutically relevant protein family, our study provides a resource for interpreting clinical resistance variants, benchmarking inhibitor specificity, and guiding the design of next-generation molecules. As precision oncology continues to advance, coupling systematic mutagenesis with rational drug development may accelerate the creation of therapies that are not only potent but also resilient to resistance.

## Methods

### Library design

To design base editing tiling libraries, we retrieved the Ensembl Canonical transcript IDs for each of the nine CDKs we chose to include in our screen. Then, using the gffutils package (v0.13) in combination with GENCODE annotations (release 44), we selected the coding sequence (CDS) for each transcript. Next, we identified all available “NG” PAM sequences on both the plus and minus strand of each gene within a window spanning the CDS as well as a 20 nucleotide (nt) buffer sequence on each end of every section of the CDS. These PAM sequences were used to generate gRNA sequences in “G+19” format, where the total protospacer length is 20 nt with the first base (position +1) substituted with a “G” to initiate U6-driven transcription. To generate final oligonucleotide designs, these gRNA sequences were inserted prior to a “F+E” optimized gRNA scaffold sequence (*94*), followed by 7 nt polyT U6 terminator sequence.

In addition, a 42-nucleotide sensor sequence was included within the oligonucleotide design **(Figure 1c)**. This sensor is a copy of the endogenous target site, and includes the 22-nt PAM sequence (2 nt) and protospacer (20 nt), as well as 10 nt of flanking sequence on either side of the PAM and protospacer, matching the sequence present at the endogenous site. This sensor is included in reverse complement orientation relative to the gRNA sequence, because we and others have shown that this decreases the rate of recombination during deconvolution (*45*). In addition, a 13-nt barcode sequence was included following the sensor sequence to allow for facile identification of the gRNA-sensor cassette.

To generate the final oligonucleotide sequences, a 5’ adapter sequence containing an Esp3I (BsmBI) site (5’-GCGTACACGTCTCACACC-3’) and a 3’ adapter sequence containing an EcoRI site (5’-GAATTCTAGATCCGGTCGTCAAC-3’) were included on either end of the oligonucleotide. Additionally, to enable subpooling and individual amplification of subsets of the library, additional adapter sequences were included on either end of the oligonucleotide sequence. The adapter sequences for each pool were:

Subpool 1: F1 = 5’-AGGCACTTGCTCGTACGACG-3’ | R1 = 5’-TTAAGGTGCCGGGCCCACAT-3’

Subpool 2: F2 = 5’-GTGTAACCCGTAGGGCACCT-3’ | R2 = 5’-GTCGAAGGACTGCTCTCGAC-3’

Subpool 3: F3 = 5’-CAGCGCCAATGGGCTTTCGA-3’ | R3 = 5’-CGACAGGCTCTTAAGCGGCT-3’

In addition to gRNAs targeting the CDS, we also included a set of intron-targeting control gRNAs in each subpool. We designed at most 4 gRNAs to mutagenize random regions within each intron, targeting regions at least 30 nt away from the nearest exonic region. Intron-targeting gRNAs also included a sensor cassette in the same design format. We also included a set of non-targeting gRNAs as another set of controls, representing ∼5% of each subpool, with a random sensor sequence included to match the design of the library (*95*).

The final oligonucleotide library was filtered to exclude sequences with additional EcoRI and Esp3I restriction sites, as this would not be compatible with our library cloning. Full details and code used to generate these libraries is provided in the associated GitHub repository (see **Code Availability** section).

### Library Cloning

The pooled library was ordered from Twist Biosciences. The lyophilized library was resuspended in 100 µl of TE buffer (pH 8.0) and diluted to create 1 ng/µl stocks. We performed *n* = 12 PCR reactions for subpool 1, *n =* 20 PCR reactions for subpool 2, and *n = 8* PCR reactions for subpool 3 with NEBNext High-Fidelity 2× PCR Master Mix (cat. no. M0541S) to amplify the library at a low cycle count. The primer sequences used for each subpool correspond with the F*X*-R*X* adapter sequences listed above, where the forward primer is equivalent to the F*X* (F1, F2, F3) adapter sequence, and the reverse primer is the reverse complement of the R*X* (R1, R2, R3) adapter sequence. The PCR reactions for each subpool were pooled and purified using the Qiagen PCR purification kit following the manufacturer’s protocols, with 10 µl of 3 M Na acetate pH 5.2 added for every five volumes of PB used per one volume of PCR reaction. Each subpool was digested with *Esp*3I and *EcoRI* (NEB), pooled, and purified. Subsequently, *n* = 15 ligations were performed for each subpool using 300 ng of digested and dephosphorylated Lenti-Trono-EFS-Blast-P2A-TurboRFP backbone (Addgene cat. no. 228893) and 3 ng of digested insert with high concentration T4 DNA Ligase (NEB, cat. no. M0202M). The ligation reactions were precipitated using QuantaBio 5PRIME Phase Lock Gel tubes before being resuspended in 3 µl of EB Buffer per four precipitated reactions. These precipitated ligation reactions were electroporated into Lucigen Endura ElectroCompetent cells (cat. no. 60242-2) before being plated on LB-carbenicillin plates and incubated at 37 °C for 16 h. We scraped the plates and collected the bacteria in 250 ml of LB-ampicillin per four plates, before incubating for 2 h at 37 °C, collecting the bacteria by centrifugation, and proceeding to perform a Qiagen Maxiprep, following the manufacturer’s protocol. After cloning, the libraries were sent for Amplicon-EZ sequencing (Azenta) to assess representation, with all subpools showing high representation and low skew.

### Base Editor & Reporter Plasmids

All plasmids were generated using Gibson Assembly strategies using NEBuilder HiFi DNA Assembly Master Mix (NEB cat. no. E2621) following the manufacturer’s protocol. The lentiviral cytosine base editor (CBE) reporter (Addgene, cat. no. 136895) was kindly provided by Dr. Lukas Dow. The CBE6-NG *in vitro* transcription (IVT) template was generated using SpCas9-CBE6b-IVT (Addgene, cat. No. 215834) and replacing the Cas9 sequence with one derived from pLenti-FNLSNG-P2A-Puro (Addgene, cat. No. 136900).

### Individual gRNA Cloning

We cloned annealed and phosphorylated oligos with *Esp*3I/*Bsm*BI-compatible overhangs encoding gRNAs into *Esp*3I/*Bsm*BI-linearized pUSEBB (*43*) plasmid backbone using high-concentration T4 DNA ligase (NEB). Plasmid sequences were confirmed via U6 sanger sequencing (Azenta) and/or whole-plasmid sequencing (Plasmidsaurus). All individual gRNA sequences are provided in **Supplementary Table 6.**

### Lentivirus production & titering

Lentiviruses were produced by co-transfection of HEK293T cells with the relevant lentiviral transfer vector and packaging vectors psPAX2 (Addgene, catalog no. 12260) and pMD2.G (Addgene, catalog no. 12259) using Lipofectamine 2000 (Invitrogen, catalog no. 11668030). Viral supernatants were collected at 48- and 72-hours post-transfection and stored at -80°C.

All viral titering was performed by plating 2 million cells per well in 6-well plates in 2 mL of media and adding freshly thawed virus of varying volumes to each well. Cells were lifted and transferred to 10-cm plates after 24-hours, and then the transduction efficiency of each well was measured with flow cytometry via fluorophore positivity quantification at 72-hours post-transduction. Library transductions were performed in the same format to ensure matching MOI.

### Cell lines

A549 cells were received from the Koch Institute ES Cell Core facility and were mycoplasma negative. These cells were cultured in F-12K media (Gibco, catalog no. 21127030) supplemented with 10% FBS and 1X Penicillin-Streptomycin (ThermoFisher).

To generate ABE- and CBE-expressing cell lines, A549 cells were transduced with EF1α-ABE8e-NG-P2A-Puromycin or EF1α-CBE6-NG-P2A-Puromycin lentivirus, respectively. Subsequently, cells were selected with 5 µg/mL of puromycin. To validate base editing activity in these cell lines, we transduced lentiviral reporter constructs wherein a defective copy of GFP was repaired by either ABE or CBE activity (*96*). We observed robust editing activity reaching nearly 100% in both cell lines.

Luciferase+ Murine *Bcr-Abl*-driven mouse acute B-cell lymphoblastic leukemia cells (*83*) were cultured in RPMI with L-glutamine (Corning, 10-040-CM), supplemented with 10% fetal bovine serum (FBS), GlutaMAX (Gibco), and 2-mercaptoethanol to a final concentration of 0.05 mM (Gibco, 21985023).

### Screening protocol

For each library subpool, transductions were performed to achieve ≥10,000X representation at an MOI of ∼0.3 for each replicate. Briefly, for each replicate, A549-CBE6-NG or A549-ABE8e-NG cells were plated with 2 millions cells/well in 6-well plates. Lentivirus encoding a given library subpool was added to each well to achieve an MOI of ∼0.3. After 24-hours, cells were lifted, each replicate’s collection of 6-well plates was combined, and cells were replated in an appropriate number of 15-cm plates containing 5 µg/mL of puromycin and 10 µg/mL of blasticidin S. In addition, a small split from each replicate was maintained without blasticidin to empirically check the MOI of each replicate. We confirmed that the MOI of each replicate was ∼0.3. Cells were passaged in media containing puromycin and blasticidin as needed until day 7 post-transduction. At this point, a ≥1,000X representation cell pellet was taken from each replicate for gDNA extraction, and each replicate was split into drug-treatment conditions or a DMSO vehicle control, with each replicate containing ≥1,000X representation. Cells from each replicate were passaged every 3 days, with a cell count taken at each split. If necessary, the dose of each drug was adjusted to maintain a representation of ≥1,000X **(Supplementary Figure 1)**. At the final timepoint of 21 days of treatment, a ≥1,000X cell pellet was taken from each replicate of each condition for subsequent gDNA extraction and library deconvolution.

For the iterative screening approach, the same protocol was followed as above. However, instead of selection with blasticidin, transduced cells were selected based on RFP positivity via flow cytometry.

For the subpool 1 screen, cells treated with SY-5609 did not survive. As a result, this condition was repeated with lower doses of SY-5609 using cryopreserved cells from day 7 post-transduction–these cells were preserved at a representation of ≥5,000X.

### Genomic DNA extraction

Genomic DNA (gDNA) was extracted from cells using the DNeasy Blood and Tissue Kit (Qiagen) following the manufacturer’s instructions. The resulting gDNA was resuspended in 100 µl of Buffer AE (10 mM Tris-Cl; 0.5 mM EDTA; pH 9.0).

### Library deconvolution

Library deconvolution was performed with two sequential PCR steps to amplify the base editing sensor construct from gDNA and add adapter sequences from next-generation sequencing. First, 8x PCRs were performed for each sample, with 25 µl of Q5 High-Fidelity 2X Master Mix (NEB#M0429S), 2.5 uL of 10 µM aliquots for the following forward and reverse primers:

5’-CGCTCTTCCGATCTCTAGCGTTCGAGTTAGGAATT-3’

5’-CTGAACCGCTCTTCCGATCTTTGTGGAAAGGACGAAACACC-3’

, along with 10 uL of molecular biology grade water, and 10 uL of gDNA. The PCR reaction was performed with the following cycle conditions: 98°C x 30sec, [98°C x 10 sec, 64°C x 30 sec, 72°C x 30 sec] for 25 cycles, 72°C x 2 minutes, 4°C hold. Next, up to 4 PCR reactions were pooled and PCR purified using the Qiagen PCR purification kit following the manufacturer’s protocols and eluted in 50 uL of Buffer EB (10 mM Tris-Cl, pH 8.5). Following PCR purification, each sample was run on a gel to isolate the appropriate sized band (245 nt), and then gel purified (Qiagen Gel Extraction Kit) and eluted in 30 uL of Buffer EB. Another round of PCR was then performed using the PCR1 product as a template. For this PCR, 2x PCRs were performed for each sample, with 25 µl of Q5 High-Fidelity 2X Master Mix (NEB#M0429S), 1 uL of 10 µM aliquots for the following forward and barcoded reverse primers:

5’-AATGATACGGCGACCACCGAGATCTACACCGCTCTTCCGATCTCTAGCGT-3’

5’-CAAGCAGAAGACGGCATACGAGAT**NNNNNNNN**CCTGCTGAACCGCTCTTCCGATCT-3’

(N represents barcode sequence)

, along with 18 uL of molecular biology grade water, and 5 uL of PCR1 template DNA. The PCR reaction was performed with the following cycle conditions: 98°C x 2 minutes, [98°C x 10 sec, 70°C x 30 sec, 72°C x 30 sec] for 10 cycles, 72°C x 2 minutes, 4°C hold. Next, the 2 PCR reactions were pooled and PCR purified using the Qiagen PCR purification kit following the manufacturer’s protocols and eluted in 50 uL of Buffer EB (10 mM Tris-Cl, pH 8.5). Following PCR purification, each sample was run on a gel to isolate the appropriate sized band (245 nt), and then gel purified (Qiagen Gel Extraction Kit) and eluted in 30 uL of Buffer EB. Following PCR purification, each sample was run on a gel to isolate the appropriate sized band (310 nt), and then gel purified and eluted in 30 uL of Buffer EB. The concentration of each purified PCR2 reaction was measured with Qubit, and then samples were pooled at equimolar concentrations prior to submission for sequencing.

### Next-generation sequencing (NGS) of library amplicons

All next-generation sequencing was performed with the Illumina NovaSeq SP100 or the Element AVITI (2x75) sequencer. All sequencing was performed with a custom primer set specifically designed for the base editing sensor construct:

R1: 5’-ACCGCTCTTCCGATCTCTAGCGTTCGAGTTAGGAATTC-3’

R2: 5’-CCTGCTGAACCGCTCTTCCGATCTTTGTGGAAAGGACGAAACACCG-3’

Index1: 5’-CGGTGTTTCGTCCTTTCCACAAAGATCGGAAGAGCGGTTCAGCAGG-3’

### Processing of sequencing data

For each sequencing run, the resulting sequencing data were demultiplexed into separate fastq files based on the sample barcode included in the PCR2 sequencing primers. We then used a custom analysis script (see associated Github repository) to filter reads with an average Phred quality score below 30. To generate count tables of gRNAs for each sample, we counted the presence of each 13 nt gRNA-sensor barcode in R1, and checked for the presence of the matching 19 nt protospacer sequence in R2, allowing 2 mismatches in the protospacer sequence.

Subsequently, to facilitate sensor editing analysis, R1 sensor reads, excluding the barcode sequence, were separated into distinct fastq files for each gRNA for each sample. We then used Crispresso2 (*97*) to quantify the editing outcomes in each set of sensor sequences, using a random sequence as the “edited” sequence such that all sequences aligned to the reference sequence with the following command: “--amplicon_seq {wt} -- expected_hdr_amplicon_seq {edited} --guide_seq {protospacer} --quantification_window_coordinates 12-32,12-32 --suppress_report --suppress_plots”. Next, using a custom analysis script, we used the Allele_frequency_table generated by Crispresso2 to determine the protein coding effects of each edit by substituting the 20 nt edited protospacer sequence into the transcript for each gene, followed by the simulation of transcription and translation, and determination of the resulting mutation(s).

All sequencing analysis scripts are provided in the associated GitHub repository.

### RNA extraction, sequencing & analysis

To isolate RNA, cell pellets from each condition and replicate were snap frozen and stored at -80°C. RNA extraction was performed using the RNeasy micro kit (Qiagen) following the manufacturer’s protocol. Next, RNA was prepared for NGS with the NEB Ultra II RNAseq Library Prep Kit for Illumina. Prepared libraries were sequenced using the Element AVITI sequencer (2x150).

To process RNA sequencing data, we used NextFlow/NF-core pipeline to filter and map reads and generate count tables. Finally, we used DESeq2 to normalize sequencing depth, quantify log_2_ fold-change, and determine adjusted p-values (*98*).

### Dose-response curves (MTT assays)

To perform dose-response curves, 2,500 A549 cells were plated in 100 uL of media in 96-well plates. After allowing cells to adhere overnight, the appropriate amount of drug was added to each well with ≥3 replicates per condition. After 72-hours, using the Cell Proliferation Kit I (MTT) from Roche (cat. No. 11465007001), assays were terminated following the manufacturer’s protocol. After allowing solubilization of the formazan crystals to occur overnight, absorbance was measured using a Tecan plate reader, with the absorbance wavelength set to 570 nm and reference wavelength set to 660 nm.

The resulting readings were normalized to the average of the vehicle treatment condition to determine the viability in each well, with the average background absorbance subtracted. These readings were then fit to a four-parameter log-logistic function using the scipy.optimize.curve_fit() function, with the maximum viability bounded between 85-100% and the minimum viability bounded between 0-100%.

### Sourcing of CDK-targeting compounds

KI-CDK9d-32 and KI-CDK9d-32N were provided by the Koehler lab. All other compounds were ordered from MedChemExpress (MCE) or SelleckChem (Selleck), with the following catalogue numbers: KB-0742 (MCE cat. no. HY-137478A), SY-5609 (MCE cat. no. HY-138293), Senexin B (MCE cat. no. HY-101800), SEL120 (Selleck cat. no. S8840), BSJ-4-116 (MCE cat. no. HY-139039), CDK12-IN-2, (MCE cat. no. HY-112626), HQ461 (MCE cat. no. HY-144981), Ribociclib (MCE cat. no. HY-15777), Palbociclib (MCE cat. no. HY-50767A), Abemaciclib (MCE cat. no. HY-16297), Atirmociclib (Selleck cat. no. E1495), Tagtociclib (MCE cat. no. HY-137894A), and INX-315 (MCE cat. no. HY-162001).

### Generation of log_2_ fold-change (LFC) and FDR values for gRNAs

To calculate log_2_ fold-change (LFC) values for each gRNA in each condition, we first normalized for the sequencing depth in each sample by calculating the reads per million (RPM) value for each gRNA. Next, we calculated the LFC of each replicate relative to the median RPM value of the corresponding base value (comparison condition), with a pseudocount of 1 added. For each sample, we generated LFC values in comparison to the DMSO-treated vehicle control, the T0 timepoint (7-days post-transduction), and the Plasmid Library.

To compute FDR values for each gRNA, we then computed an empirical null distribution of LFC values in each condition by determining the LFC of non-targeting control gRNAs in each replicate. To generate a sufficiently large null distribution for p-value calculations, we merged the LFC values of control gRNAs across replicates in a given condition, and then performed bootstrapping, sampling with replacement to generate a null distribution of 10,000 LFC values. We then used these empirical null distributions, which capture the technical and biological variability in each condition, to calculate empirical p-values for each gRNA in each replicate. These p-values were simply defined as the minimum of (p_high_, p_low_), where p_high_ is defined as the 1 + the number of gRNAs in the empirical null distribution with a LFC value greater than or equal to the observed LFC value of the gRNA under consideration divided by 1 + the size of the empirical null distribution. Similarly, p_low_ is similarly defined as 1 + the number of gRNAs in the empirical null distribution with a LFC value less than or equal to the observed LFC value of the gRNA under consideration divided by 1 + the size of the empirical null distribution. In other words, the empirical p-values are defined as the percentile rank of the observed LFC value relative to the empirical null distribution.

We then combined p-values across replicates in a given condition using Fisher’s method (Fisher’s combined probability test) with the combine_pvalues() function from the scipy.stats package. We then computed FDR values using the Benjamini-Hochberg method, again using the scipy.stats package, with the multipletests() function. These FDR values were used for calling significantly enriched or depleted gRNAs in downstream analysis.

The code used to analyze sequencing data is provided in the associated GitHub repository. The results of this analysis are provided in Supplementary Tables.

### Quantification of sensor editing

After quantifying sensor editing in each sample using Crispresso2 combined with a custom analysis pipeline to determine the protein coding effects of each edit (see “Processing of sequencing data” section above), we merged together conditions to generate a maximum likelihood estimate (MLE) of sensor editing at both the DNA and amino acid level. To do so, we summed the editing outcomes across replicates and divided by the total number of sensor reads across replicates. For example, if we observed 10 reads out of 100 with a given edit in one replicate, and 50 reads out of 200 with the same edit in another replicate, these would be merged to generate an MLE of (10+50)/(100+200) = 20% editing for that particular edit. This MLE calculation was used to determine the editing rate at both the DNA level, as well as the amino acid level, given that distinct mutations at the DNA level can produce the same amino acid substitutions. The MLE was calculated for each condition, and these results are provided in the associated GitHub repository. For all analyses, we merged across all conditions to maximize the sample size of profiled, noting a high correlation in sensor editing among the different samples.

The above analyses provide tables of edits at the DNA and amino acid level, including compound mutations. To enable more facile structure-function analysis, we further decomposed these editing tables to quantify the frequency of every single amino acid variant (SAV) produced by each gRNA. For example, if sensor editing indicated that a given gRNA was producing a M42G edit at a frequency of 25% and a M42G_H43R mutation at a frequency of 40%, the resulting SAV frequencies would be: M42G = 65% and H43R = 40%. These SAV frequencies were used in the structure-function analysis presented in Figure 5 & 6, and to generate logo plots of sensor editing.

For comparison with simulated editing in the +4 to +8 region of the protospacer **(Figure 1)**, we simulated the mutagenesis of every base-editing amenable nucleotide in the +4 to +8 region of the coding sequence targeted by each gRNA. We generated each combination of possible edits produced by each guide and used these sequences to predict the editing at the amino acid level.

All sensor analysis and quantification methods are included in the associated GitHub repository.

### Variant Effect Prediction analysis with dbNSFP

We used dbNSFP to compute Variant Effect Prediction (VEP) scores for amino acid variants present in sensor editing data (*99*). Of note, dbNFSP is only compatible with SNPs, so the quantification was only run on the amino acid variants that could be generated with a single nucleotide change (the majority of the observed SAVs).

### KLIFS analysis

KLIFS annotations for each CDK were accessed from the missense kinase toolkit (https://mkt-app.streamlit.app/) (*100*) and the resulting json files containing annotation information were processed to produce a centralized table containing the KLIFS indexing for each CDK.

### Multiple Sequence Alignment (MSA)

Multiple sequence alignment was performed using Clustal Omega on the protein sequences of each gene (*101*). Custom analysis scripts were then used to process the MSA to generate a unified indexing to enable protein comparison (see associated GitHub repository).

### Structural Analysis

All structural visualizations were generated using ChimeraX (v1.1.0.1) (*102*). The “matchmaker” command was used to align structures to provide a consistent view for comparison of different structures. The “mutationscores” command was used to generate structures colored by residue essentiality **(Figure 5c)**. All PDB and Alphafold codes are provided in the associated figure legends.

### Flow Cytometry

Single cell suspensions were prepared from cells after collection by passing specimens through a 100-µM cell strainer. Cells were resuspended in FACS buffer (PBS 1X + 2.5% fetal bovine serum). For all experiments, flow cytometric analysis was conducted using either BD FACS Symphony A1 or Celesta analyzers. Fluorescence-assisted cell sorting was performed in either BD FACS Aria II or Sony MA900 cell sorters. Data were analyzed using FlowJo v10.10, as indicated in the figure legends.

### *In vitro* transcription

The IVT ABE template was linearized with BbsI and purified via phenol chloroform extraction and ethanol precipitation. The CBE IVT template was linearized using a PCR protocol detailed in Doman et al (*103*) and purified using the QIAquick PCR purification kit (Qiagen cat. no. 28104) following the manufacturer’s protocol. All IVT reactions were performed on the linearized template for 2 hours @ 37 °C using N1-methylpseudouridine-5’-triphosphate and cleanCap AG co-transcriptional capping (TriLink cat. no. N-1081-10 and N-7113-10). DNA template was then digested away with TURBO DNase (Thermo cat. no. EN0521). mRNA was then purified using lithium chloride precipitation. RNA was quantified and QC was performed by running the RNA on a glyoxal gel to confirm the appropriate size and lack of significant RNA degradation (NorthernMax AM8678). Base editor mRNA was also tested for functionality by nucleofection into reporter cell lines.

### Nucleofection

All electroporations were performed using a Neon Transfection System (Invitrogen, discontinued) or a Neon NxT Electroporation System (Invitrogen, cat. no. NEON1SK). For all screens and reporter activity assays, 500,000 cells were electroporated per reaction with the following parameters: 1600V, 20 ms, 1 pulse. Electroporations used either 10 ug (Cas9, ABE) or 20 ug (CBE) of mRNA per reaction.

### Ethics statement

The research conducted in this study was conducted ethically and complies with all relevant guidelines and regulations. Animal studies were performed under strict compliance with the Committee of Animal Care (CAC) at the Massachusetts Institute of Technology (#2203000315).

### Animal housing and maintenance

All mouse experiments were conducted under Institutional Animal Care and Use Committee (IACUC)-approved animal protocols at the Massachusetts Institute of Technology (MIT). The mouse strains used in this study included male C57BL/6J mice (The Jackson Laboratories), as indicated in the figure legends. All experimental mice used were 8-12 weeks old. Mice were housed under social conditions (two to five mice per cage) on a 12- hour dark/12-hour light cycle, ambient temperature 21 °C ± 1 °C, and humidity 50% ± 10%. All animals were housed in the pathogen-free animal facility of the MIT Koch Institute, in accordance with the animal care standards of the institutions. Food and water was provided *ad libitum*. All animal research at MIT is conducted under humane conditions with utmost regard for animal welfare. The animal care facility staff is headed by a chief veterinarian and includes a veterinary assistant, animal care technicians, and administrative support. MIT adheres to institutional standards for the humane use and care of animals, which have been established to assure compliance with all applicable federal and state regulations for the purchase, transportation, housing, and research use of animals.

### Animal studies and treatment

All mouse experiments were approved by the MIT Committee on Animal Care (CAC) and the Department of Comparative Medicine (DCM). B-ALL cells were prepared for transplantation by resuspension in 100-200uL PBS (Corning, 21-031-CV) and loaded in 27.5-gauge syringes (Becton Dickinson, Catalog# BD 305620). All cell solutions were administered by tail vein injection. C57BL/6J mice were injected intravenously with 1-2.0x10^6^ cells. All mice were closely monitored and euthanized upon the appearance of morbidity, in accordance with CAC and DCM policy. The size of each animal cohort was determined by estimating biologically relevant effect sizes between conditions and using the minimum number of animals that could reveal statistical significance using the indicated tests of significance.

For Cdk inhibition experiments, mice were treated with either KI-CDK9d-32 (2mg/kg) or SY-5609 (2mg/kg) 5 days after transplantation. Tumor burden was monitored over time by bioluminescence imaging, as indicated in the figure legends. For quantification of leukemia burden, mice were euthanized 10 days after transplantation and spleens were harvested. Single cell suspensions were generated from spleens after mechanical dissociation of tissue and lysis of red blood cells (Red Blood Cell Lysis Buffer-HybriMax; Sigma-Aldrich (R7757)). Flow cytometry was utilized to identify BFP+ B-ALL cells to calculate splenic leukemia burden, as indicated in figure legends. For survival experiments, mice were monitored for lethargy, weight loss, and/or limb paralysis, at which point they were euthanized.

### Bioluminescence studies

XenoLight D-Luciferin Potassium Salt D (PerkinElmer, Catalog# 122799) was used as substrate for standard bioluminescent imaging (30 mg/mL in PBS). Luciferin was administered via intraperitoneal injection at a dose of 165 mg/kg. Mice were anesthetized with 2.5% isoflurane (Isospire Inhalation Anesthetic, #ISO-250), delivered at 1 L per minute in O2. Animals were imaged over a 6-minute timecourse on a Xenogen IVIS system at consistent exposures between groups with small binning. Data were analyzed using Living Image software (Version 4.8.2; Revvity).

## Data Availability

Raw sequencing will be deposited in the SRA. All other processed datasets and source data used to generate figures is provided in the associated <u>GitHub repository</u>.

## Code Availability

All analysis scripts, as well as Jupyter notebooks for generating each figure that appears in the paper, are available at the following GitHub repository: https://github.com/samgould2/CDK-BE-mutagenesis.

## Supplementary Tables

- **Supplementary Table 1**: CDK base editing tiling library
- **Supplementary Table 2**: Editing information for ABE screens
- **Supplementary Table 3**: Editing information for CBE screens
- **Supplementary Table 4:** LFC/FDR information for ABE screens
- **Supplementary Table 5**: LFC/FDR information for CBE screens
- **Supplementary Table 6:** Sequences of individually tested gRNAs

## Acknowledgments

We thank all members of the Sánchez-Rivera laboratory for feedback and support. We also thank Michael Yaffe, Pau Creixell, Lukas Dow, Alberto Ciccia, Yu-Jui Ho, John Chodera, Sukrit Singh, Iván Pulido, Jonathan Weissman, and Yadira Soto-Feliciano for scientific discussions and overall support. We thank Claire Glickman for laboratory management support and Paul Thompson for administrative support. We also thank Aurora Burds Connor and Wontaek Chung for mouse husbandry and experimental assistance. We acknowledge and thank the Koch Institute’s Robert A. Swanson (1969) Biotechnology Center for technical support, especially the Barbara K. Ostrom (1978) Bioinformatics Facility. Work in the Sánchez-Rivera laboratory is supported by the Howard Hughes Medical Institute (Hanna Gray Fellowship, GT15656), the V Foundation for Cancer Research (V2022–028), the National Cancer Institute (Cancer Center Support Grant P30-CA014051, NCI 1P01CA291694–01A1), the Virginia and D.K. Ludwig Fund for Cancer Research, the MIT Center for Precision Cancer Medicine (CPCM), the MIT HEALS Initiative, the Koch Institute Frontier Research Program, the Casey and Family Foundation Cancer Research Fund, Michael (1957) and Inara Erdei Fund, the MIT Research Support Committee, the Upstage Lung Cancer Foundation, and a Traditional Project Award from the Bridge Project, a partnership between the Koch Institute for Integrative Cancer Research at MIT and the Dana-Farber/ Harvard Cancer Center. S.I.G., G.A.J., and M.E.C. were supported by NIH T32GM136540. S.I.G. was also supported by a MIT School of Science Fellowship in Cancer Research. G.A.J. was also supported by a Margaret A. Cunningham Immune Mechanisms in Cancer Research Fellowship Award. S.C. is supported by NCI P30CA008748, R35CA305347, and the BCRF. We also thank the Koch Institute Swanson Biotechnology Center for technical support, especially the Flow Cytometry Core and the Barbara K. Ostrom (1978) Bioinformatics Facility and the Genomics Facility.

## Author Contributions Statement

S.I.G., and F.J.S.R. conceived the project and wrote the manuscript. S.I.G. performed all analyses of experimental data. S.I.G., J.A., A.J., G.A.J., M.E.C., A.N.W., and E.B. performed experiments. L.B.B. and S.C. performed analysis of clinical data and provided comments on the manuscript. M.A.T., Y.X., Z.K., M.T.H. and A.N.K. provided technical and conceptual advice and experimental reagents. F.J.S.R. supervised the work and secured funding.

---

**Supplementary Figure 1.**
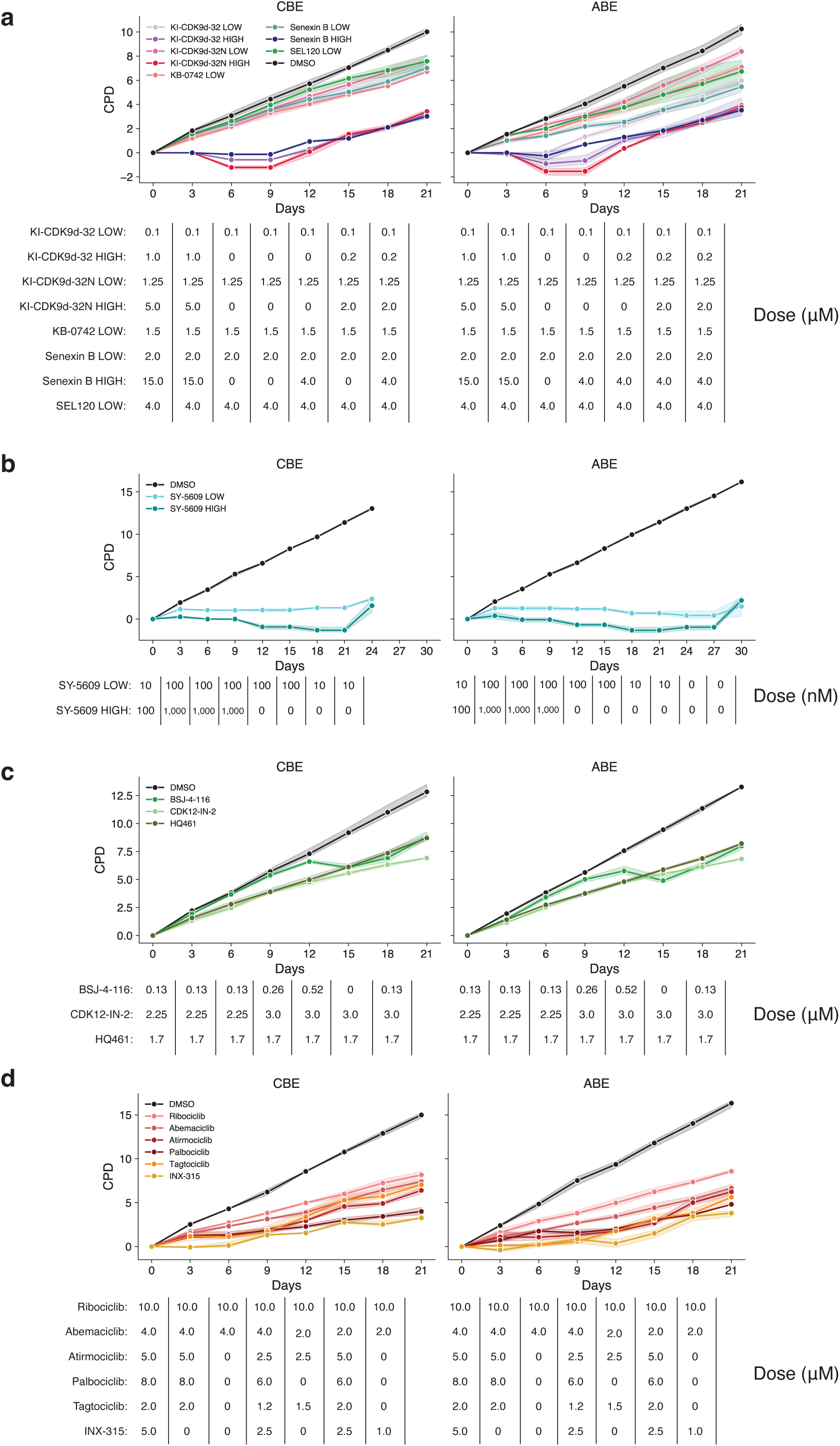
Cumulative population doublings in each screen. Cumulative population doublings (CPD), defined as the sum of log_2_(cell count/cells plated) for **(a)** the subpool 1 screen, **(b)** the re-screen of subpool 1 in the presence of SY-5609, **(c)** the subpool 2 screen, and **(d)** the subpool 3 screen. Points represent the average CPD value across three replicates, with the shaded region indicating the 95% confidence interval of the mean value. Drug doses at each timepoint, which had to be adjusted for certain conditions based on cell count and morphology, are indicated below each plot.

**Supplementary Figure 2.**
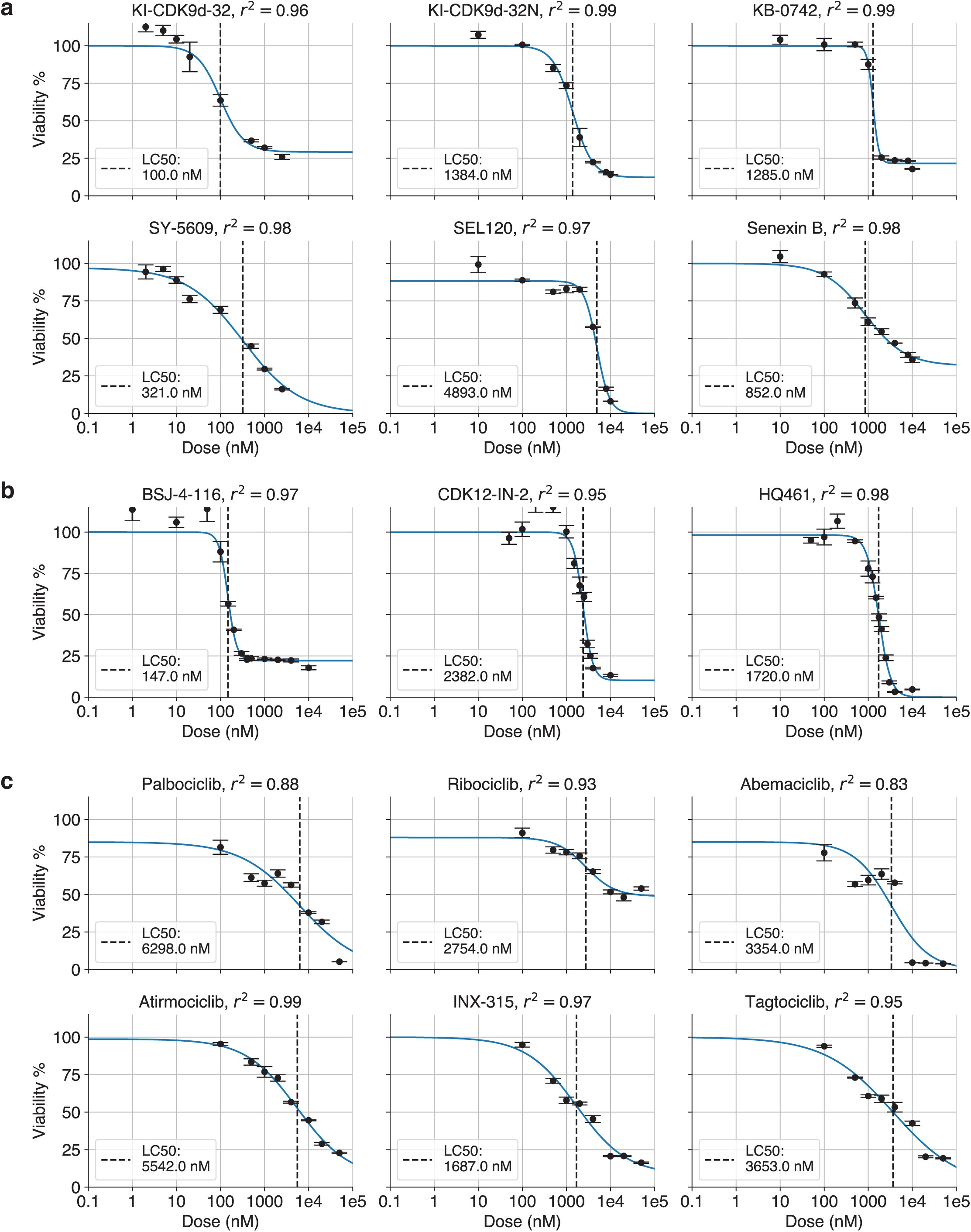
Dose-response curves for each drug. Dose-response curves for drugs used in **(a)** subpool 1, **(b)** subpool 2, and **(c)** subpool 3. Error bars at each point indicate standard deviation of three technical replicates. Curves fit with four parameter log-logistic function, with LC50 calculated as the concentration where half of the maximal cell death occurs.

**Supplementary Figure 3.**
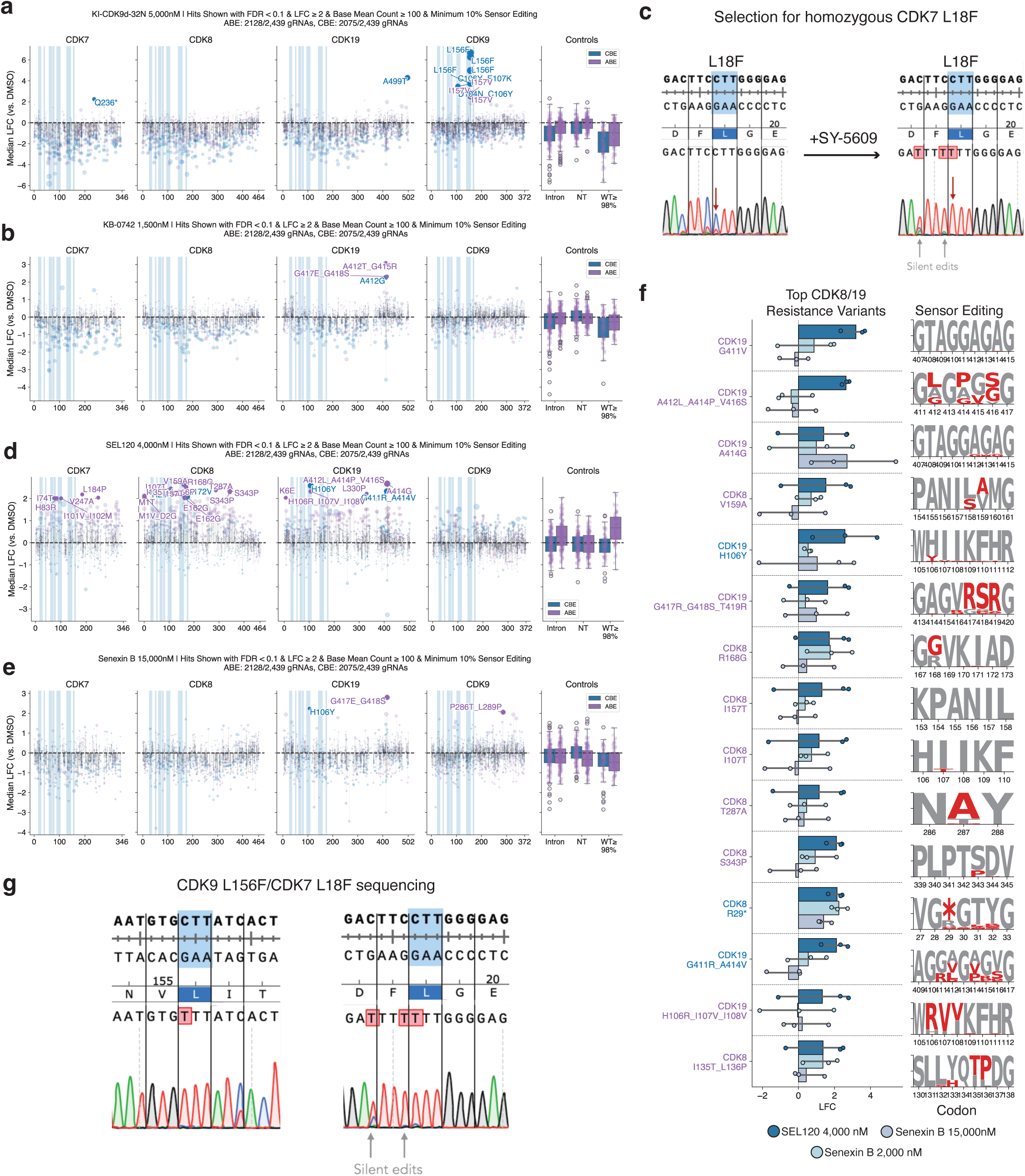
Additional information about subpool 1 and iterative screens. **a)** Scatterplot of median gRNA enrichment (log_2_ fold-change) in the presence of 5,000 nM KI-CDK9d-32N relative to DMSO-treated control. Each dot represents a gRNA, filtered to exclude gRNAs below 10% sensor editing and with a base mean count ≤ 100. Hits labelled with FDR < 0.1 & LFC ≥ 2. ABE screen results shown in purple; CBE in blue. Regions shaded blue indicate ATP binding site residues. **b)** Same as (a), but for cells treated with KB-0742 (1,500 nM). **c)** Sanger sequencing of cells transduced with CDK7 L18F gRNA before and after selection with SY-5609. **d)** Same as (a), but for cells treated with SEL120 (4,000 nM). **e)** Same as (a), but for cells treated with Senexin B (15,000 nM). **f)** Top resistance variants in CDK8/19, as ranked by log_2_ fold-change in SEL120 4,000 nM treatment condition. **g)** Sanger sequencing of CDK9 L156F/CDK7 L18F cells.

**Supplementary Figure 4.**
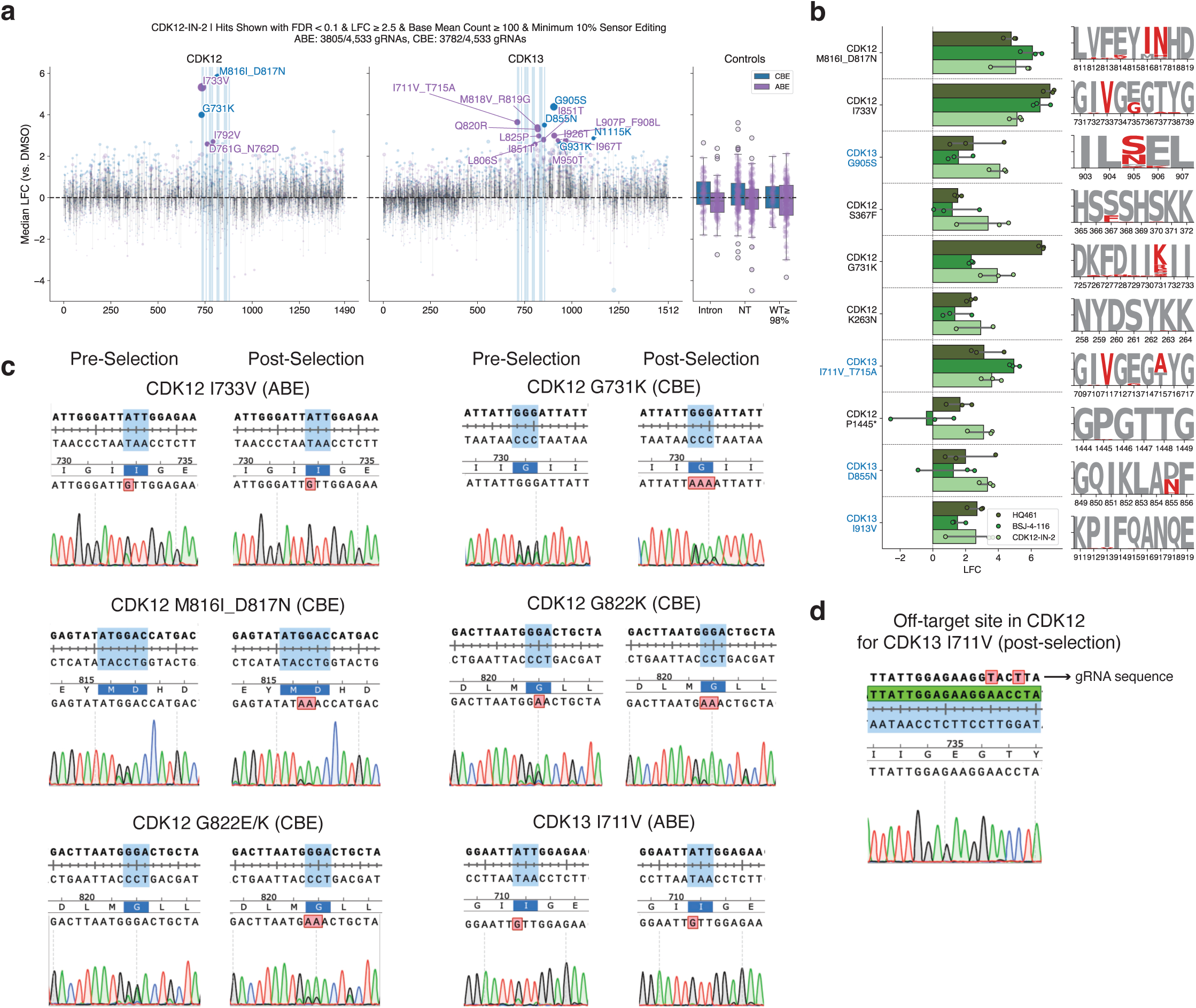
Identification and validation of resistance variants in CDK12/13. **a)** Scatterplot of median gRNA enrichment (log_2_ fold-change) in the presence of CDK12-IN-2 relative to DMSO-treated control. Each dot represents a gRNA, filtered to exclude gRNAs below 10% sensor editing and with a base mean count ≤ 100. Hits labelled with FDR < 0.1 & LFC ≥ 2. ABE screen results shown in purple; CBE in blue. Regions shaded blue indicate ATP binding site residues. **b)** Top resistance variants in CDK12/13, as ranked by log_2_ fold-change in CDK12-IN-2 treatment condition. **c)** Sanger sequencing of arrayed validation of CDK12/13 mutations before and after selection with HQ461. **d)** Sanger sequencing of CDK12 off-target site for cells transduced with CDK13 I711V gRNA and selected with HQ461, indicating no off-target editing.

**Supplementary Figure 5.**
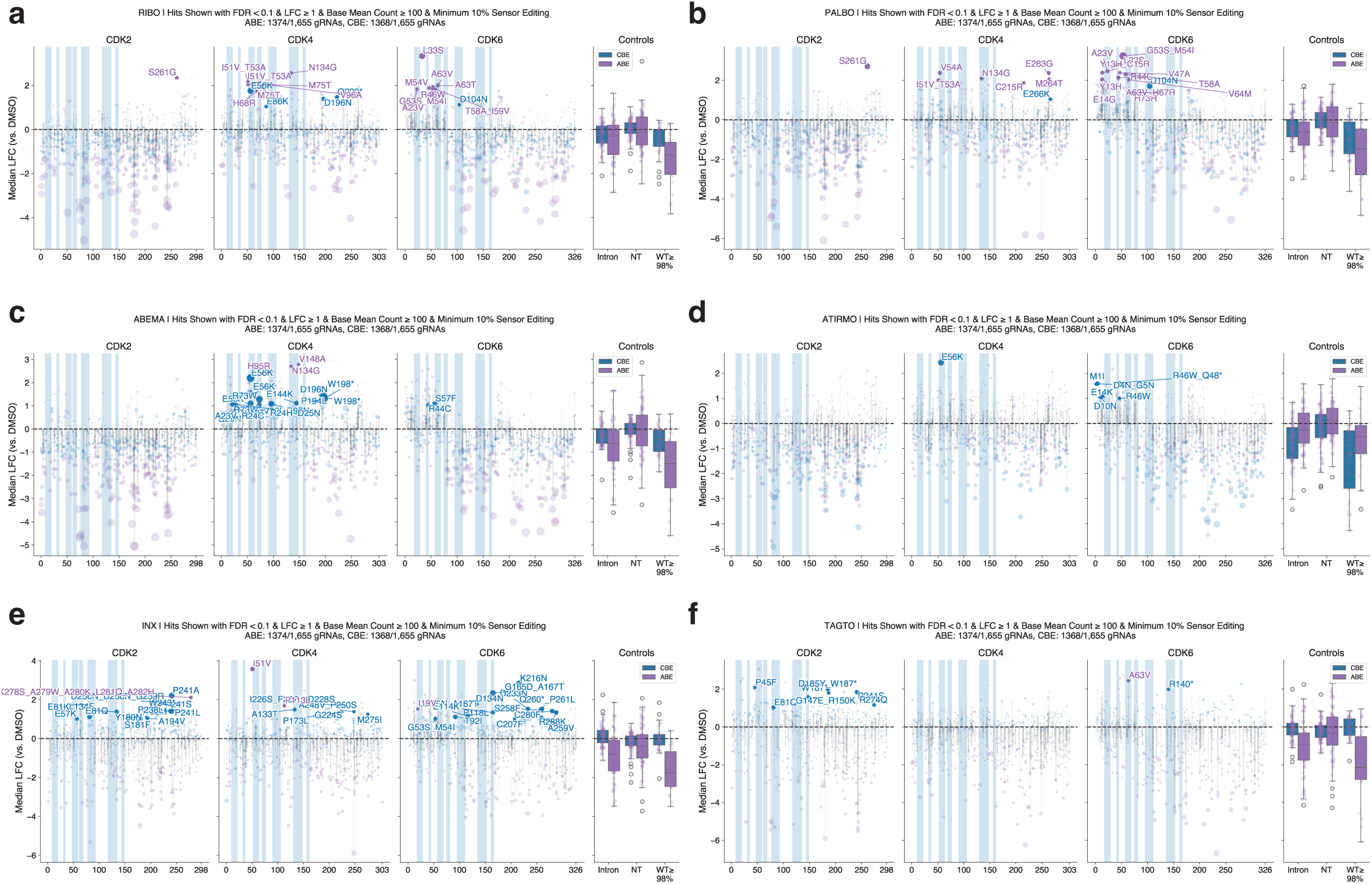
Identification of resistance variants in CDK2/4/6. Scatterplots of median gRNA enrichment (log_2_ fold-change) in the presence of **a)** Ribociclib, **b)** Palbociclib, **c)** Abemaciclib, **d)** Atirmociclib, **e)** INX-315, or **f)** Tagtociclib relative to DMSO-treated control. Each dot represents a gRNA, filtered to exclude gRNAs below 10% sensor editing and with a base mean count ≤ 100. Hits labelled with FDR < 0.1 & LFC ≥ 1. ABE screen results shown in purple; CBE in blue. Regions shaded blue indicate ATP binding site residues.

